# *Borrelia burgdorferi* infection-enhanced *Ixodes scapularis* serpins S12c5 and S13c5, target MASP1/3 to inhibit MBL-pathway complement activation and promote spirochete survival

**DOI:** 10.64898/2026.06.26.734773

**Authors:** Emily Bencosme-Cuevas, Tae K. Kim, Thu-Thuy Nguyen, Moiz A. Ansari, Shahbaz M. Khan, Payton G. Smith, Kyla R. Herr, Yungui Guo, William H. Witola, Jon T. Skare, Brandon L. Garcia, Klaudia I. Kocurek, L. Garry Adams, Stefan H.E. Kaufmann, Albert Mulenga

## Abstract

Tick serine protease inhibitors (serpins) are critical for tick feeding success and pathogen transmission. We previously demonstrated that two unique serpins, S12c5 and S13c5, are tandemly duplicated on *Ixodes scapularis* chromosome 5 and are injected into the host at heightened levels during feeding by *Borrelia burgdorferi* (Bb)-infected nymphs. Here, we characterize the biochemical properties and immunomodulatory functions of these serpins and investigate their roles in promoting Bb transmission. Despite their response to Bb infection, S12c5 and S13c5 share less than 20% overall amino acid identity and differ critically at their reactive center loop (RCL) P1 site, which determines substrate specificity: S13c5 contains a basic arginine residue, while S12c5 contains a polar serine residue. Consistent with these differences, recombinant S13c5 efficiently inhibited innate immune proteases including fXa, plasmin, and trypsin IV (SI: 2.5, 1.9, and 1.0; ka: 8.56 × 10³, 3.97 × 10⁵, and 1.08 × 10⁵ M⁻¹s⁻¹, respectively), and consequently delayed plasma clotting via the common pathway, whereas rS12c5 did not inhibit these proteases. However, despite molecular modeling predicting stronger interactions between S13c5 and complement proteases, both serpins inhibited MBL-pathway complement activation by targeting MASP1/3. Deletion of the RCL domain abolished this inhibitory activity, confirming both proteins function as typical inhibitory serpins. Functionally, rS13c5 enhanced Bb colonization in C3H mouse organs. Interestingly, immune response assays revealed a trade-off: S13c5, despite its broader inhibitory activity, was less immunogenic than S12c5. Collectively, these findings establish S12c5 and S13c5 as Bb transmission factors that promote spirochete survival through disruption of host innate immunity.

## Introduction

Vector-borne diseases (VBDs) account for 17% of infectious diseases world-wide (1). Ticks are second to mosquitoes for transmission of most important human vector-borne pathogens (1). In 2017, the WHO Assembly adopted a 13 yearlong (2017–2030) resolution calling Member States to develop and strengthen vector control strategies (2). Shortly after in 2019, the Kay Hagan Tick Act was signed into law in the United States of America (USA), requiring the Department of Health and Human Services to develop a strategy addressing VBDs including those that are tick-borne.

In the United States, Lyme disease (LD), caused by *Ixodes scapularis*-transmitted *Borrelia burgdorferi* (Bb) is the most common human VBD. According to the Centers for Disease Control and Prevention (CDC), more than 30,000 confirmed cases and approximately 476,000 suspected cases occur annually in humans (3, 4). There are ongoing efforts to develop vaccines for humans to prevent LD (5, 6). Despite there being several LD vaccines for use in canines (7), none are currently approved for human use (8).

In the absence of effective vaccines to prevent LD in humans, tick antigen–based vaccines are emerging as a promising alternative. This approach builds on early studies showing that hosts can develop strong anti-tick immunity after repeated infestations (9) and that the response is driven by salivary gland antigens (10). Notably, such immunity, acquired through repeated tick exposure protected hosts against Bb transmission (11–13), underscoring the importance of tick saliva components. Moreover, active immunization of mice with tick saliva proteins has been shown to reduce transmission of LD agents (14–16). Conversely, tick saliva and salivary gland extracts enhanced pathogen replication (17, 18), facilitate evasion of innate immunity ex vivo (19), and enhance organ colonization in needle-inoculated mice (20, 21). These findings highlight the significance of tick saliva transmission factors of LD agents as attractive targets for developing tick antigen-based anti-tick vaccines to prevent pathogen transmission (22–26).

Tick feeding begins with the insertion of the hypostome into the host epidermal and dermal skin layers (27), creating a wound that triggers immediate host defense mechanisms, including complement activation, inflammation, hemostasis, and pain (28). Ticks successfully counteract host defenses through secretion of several immunomodulating factors. Our lab and others have extensively identified and characterized several tick saliva proteins that are secreted during tick feeding (22, 29–38). Innate defense systems that the tick must inhibit to feed and transmit pathogens are predominantly serine protease-mediated pathways such as inflammation, blood clotting, and complement activation that are tightly regulated by serpins (39, 40). On this basis, tick saliva serpins were proposed as key modulators of host defense against ticks and thus, attractive candidate targets for tick antigen-based vaccines against tick feeding (41). Our lab and others have described serpins in several tick species (25, 26, 42, 43) and *I. scapularis* (24, 44–46). Functional analysis studies have confirmed that tick saliva serpins regulate innate immune system protease effectors (23, 33, 47–49).

Our recent study identified serpins among highly induced or secreted proteins in the saliva of Bb–infected *I. scapularis* nymphs (24), suggesting their potential role in Bb transmission. Consistent with this, we previously showed that two infection-responsive serpins, IxsS17 (47) and IxsS41 (49), inhibit innate immune proteases involved in blood clotting, inflammation, and complement activation. Building on these findings, we analyzed the updated *I. scapularis* genome (50) and identified 74 serpin-encoding genes, revising previous estimates (47, 50, 51). We also refined the classification of tick serpins based on their chromosomal organization (52). Expanding our understanding of serpins in tick physiology and pathogen transmission, this study aimed to characterize the roles of two serpins, S12c5 and S13c5, in tick feeding and Bb transmission. We focused on these two proteins as a pair because they exhibited similar secretion dynamics in Bb infected ticks (24) despite showing low amino acid identity at less than 20%. Interestingly, despite low overall amino acid identity, their genes share a similar organization on *I. scapularis* chromosome 5: each is duplicated and arranged in tandem (52). Here, we show that S12c5 and S13c5 function as inhibitors of complement activation via the lectin pathway by directly binding MASP1/3, the pathway’s key activator (53, 54). Notably, S13c5 efficiently inhibited proteases involved in blood clotting and inflammatory pathways, acted as an anticoagulant, and promoted Bb colonization of the host. Interestingly, despite being highly secreted by infected ticks, rabbits fed on by uninfected ticks generate high antibody levels to these proteins compared to rabbits fed on by infected ticks. Equally notable, we found a functional trade-off: S12c5, which has reduced protease inhibitory functions, is highly immunogenic compared to S13c5, which exhibits broad inhibitory activity. This suggests that the highly immunogenic S12c5 may serve as a decoy to protect the functionally critical S13c5 from the host’s immune response.

## Results

### S12c5 and S13c5 are unique but conserved across tick species

Serpins S12c5 and S13c5, which each occur in duplicate and are arranged in tandem on *I. scapularis* chromosome 5 (52), are highly conserved across several tick species. Pairwise comparison between S12c5 (XP_040071636.1) and S13c5 (XP_029834128.2) revealed low overall identity of 12%, and 17.2%, respectively within their functional reactive center loop (RCL) domains. When compared to other protein sequences in GenBank, S12c5 is 95% conserved in *Ixodes persulcatus* (KAG0414789.1), and 61–71% in Metastriata tick species, including *Dermacentor silvarum* (XP_049517673.1), *Rhipicephalus sanguineus* (XP_037510184.1), *Hyalomma asiaticum* (KAH6937721.1), *Dermacentor andersoni* (XP_050043950.1), *Haemaphysalis longicornis* (KAH9371121.1), and *Rhipicephalus microplus* (KAH8009656.1). Similarly, S13c5 showed 99% identity to *Ixodes persulcatus* (KAG0414788.1) and 66–68% identity to Metastriata tick serpins, including *Amblyomma americanum* (JAI08960), *Dermacentor andersoni* (XP_050043949.1), *Dermacentor silvarum* (XP_049517673.1), *H. longicornis* (KAH9371123.1), *Rhipicephalus haemaphysaloides* (QNV48485.1), *Rhipicephalus microplus* (XP_037285924.1), and *Rhipicephalus sanguineus* (XP_037510183.1). Within the RCL, S12c5 and S13c5 are 100% identical to *Ixodes persulcatus* serpins. Notably, although the overall amino acid identities of S12c5 and S13c5 to Metastriata tick serpins are below 71%, their RCLs are highly conserved up to 90% and 81%, respectively (Appendix Figs. S1A and S1B).

### Pichia pastoris expressed recombinant (r) S12c5 and S13c5 are glycosylated

Silver staining and western blotting analyses confirmed that both rS12c5 (Supplementary Fig. S2A) and rS13c5 (Supplementary Fig. S2B) were highly expressed in *Pichia pastoris* within 24 h of methanol induction and increased in abundance throughout five days of culture. Thereafter, large scale (1 liter) expression batches were cultured for five days and rS12c5 and rS13c5 were affinity purified under native conditions and purity was confirmed by silver staining of native SDS-PAGE (Supplementary Figs. S2C and S2D). Protein bands were diffuse suggesting glycosylation (40-110 kDa), which was confirmed by treatment with deglycosylation enzyme mix (DG^+^) which removes both N-and O-linked glycans (Supplementary Figs. S2C and S2D).

### rS13c5, but not rS12c5, is an efficient inhibitor of innate immune system proteases

To gain their functional insights in tick feeding regulation and pathogen transmission, inhibitory activity of affinity purified rS12c5 and rS13c5 (1 μM) was evaluated against 18 host defense system proteases in substrate hydrolysis assays. At 1 μM concentrations, rS12c5 moderately inhibited chymotrypsin (67.6%), chymase (65.4%), and cathepsin G (51.9%) while inhibition of the rest of the proteases ranged from 0-46% (Fig. 1A). In contrast, rS13c5 strongly inhibited activity of proteases associated with tissue regeneration, blood clotting, and inflammation system proteases including trypsin IV (99.9%), fXa (97.8%), plasmin (96.2%), pancreatic trypsin (96.2%), and fXla (86.6%) (Fig. 1B). Additionally, rS13c5 also moderately inhibited other inflammation system proteases, proteinase 3 (64.4%), cathepsin G (61.7%), chymase (60.9%), blood clotting fIXa (60.6%), and the remaining proteases by less than 40% (Fig. 1B).

**Figure 1.**
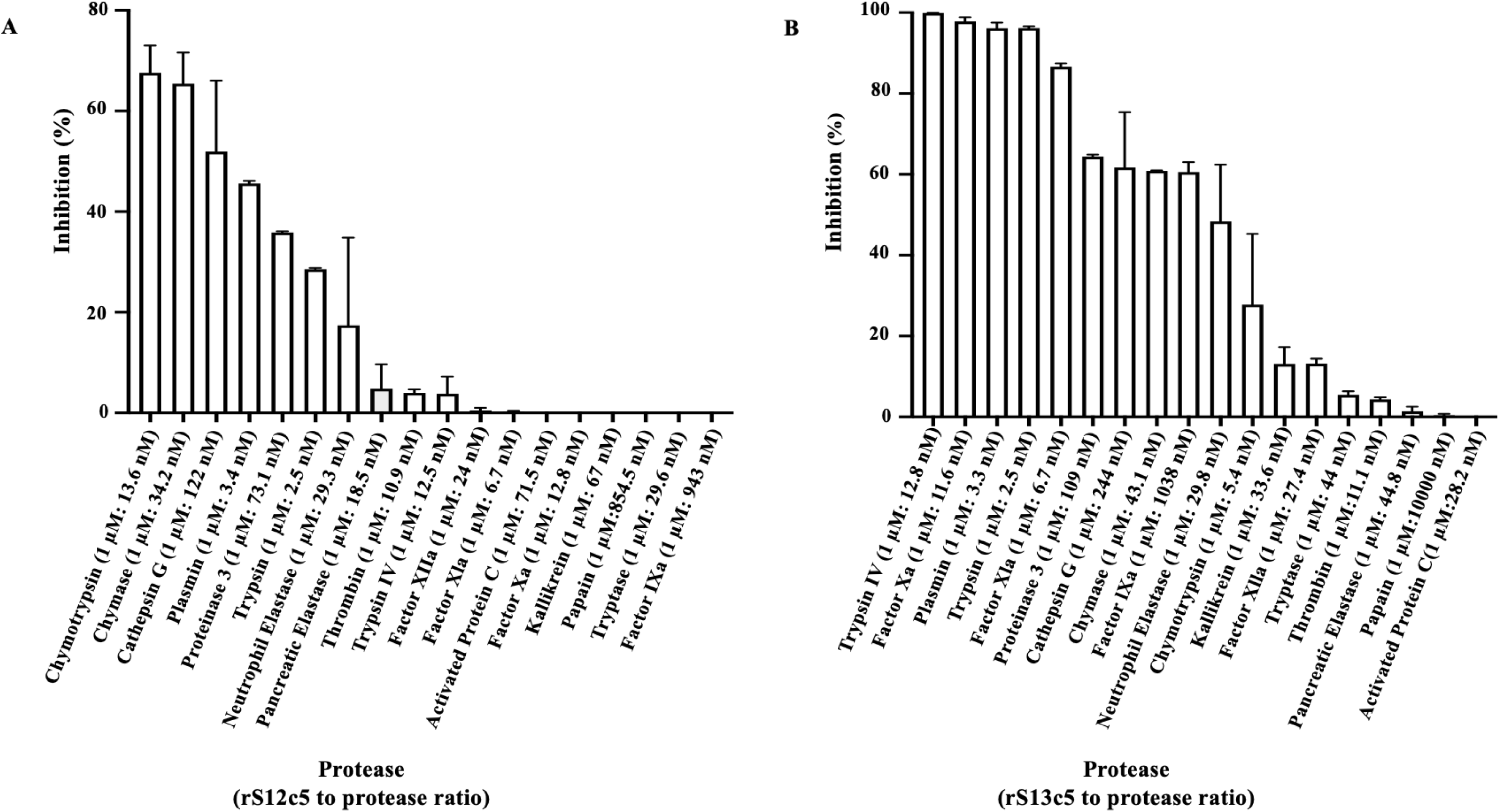
S13c5 is an inhibitor of tissue repair and blood clotting systems serine proteases and S12c5 is a moderate inhibitor. Excess amount (1 µM) of yeast expressed rS12c5 or rS13c5 was preincubated for 15 min at 37 °C with different mammalian proteases at indicated serpin-to-protease molar ratios. The chromogenic pNA substrates respective to each protease was added at a 200 µM final concentration in a 100 µL total reaction. Immediately after, substrate hydrolysis (*A*_405nm_) was recorded on the Synergy H1 plate reader every 11 s for a total for 15 min. Data were analyzed using one phase decay in PRISM 10 and calculated plateau values represented residual enzyme activity. Data are presented as percent inhibition of each protease by either rS12c5 (A) or rS13c5 ± SEM. Ratio of serpin to protease are indicated on the X axis.

We suspected that molar ratios of native S13c5 protein to its native target protease at the feeding site is likely close to 1 (55). Thus, to assess the physiological relevance of our substrate hydrolysis data, we evaluated the stoichiometry of inhibition (SI) for rS13c5 against proteases that were inhibited by more than 80%. This analysis estimated that 100% activity of a single molecule of blood clotting fXa, trypsin IV, and plasmin is inhibited by 2.5, 1.9, and 1.0 molecules of rS13c5 respectively (Fig. 2A). We next assayed the rate of the inhibitory reactions (*ka*) as described (55). We found that the rate of rS13c5 inhibition of fXa was relatively slower at 8.56 x 10^3^M^-1^s^-1^ and was faster against plasmin and trypsin IV at 3.97 x 10^5^M^-1^s^-1^ and 1.08 x 10^5^M^-1^s^-1^ respectively (Fig. 2B).

**Figure 2.**
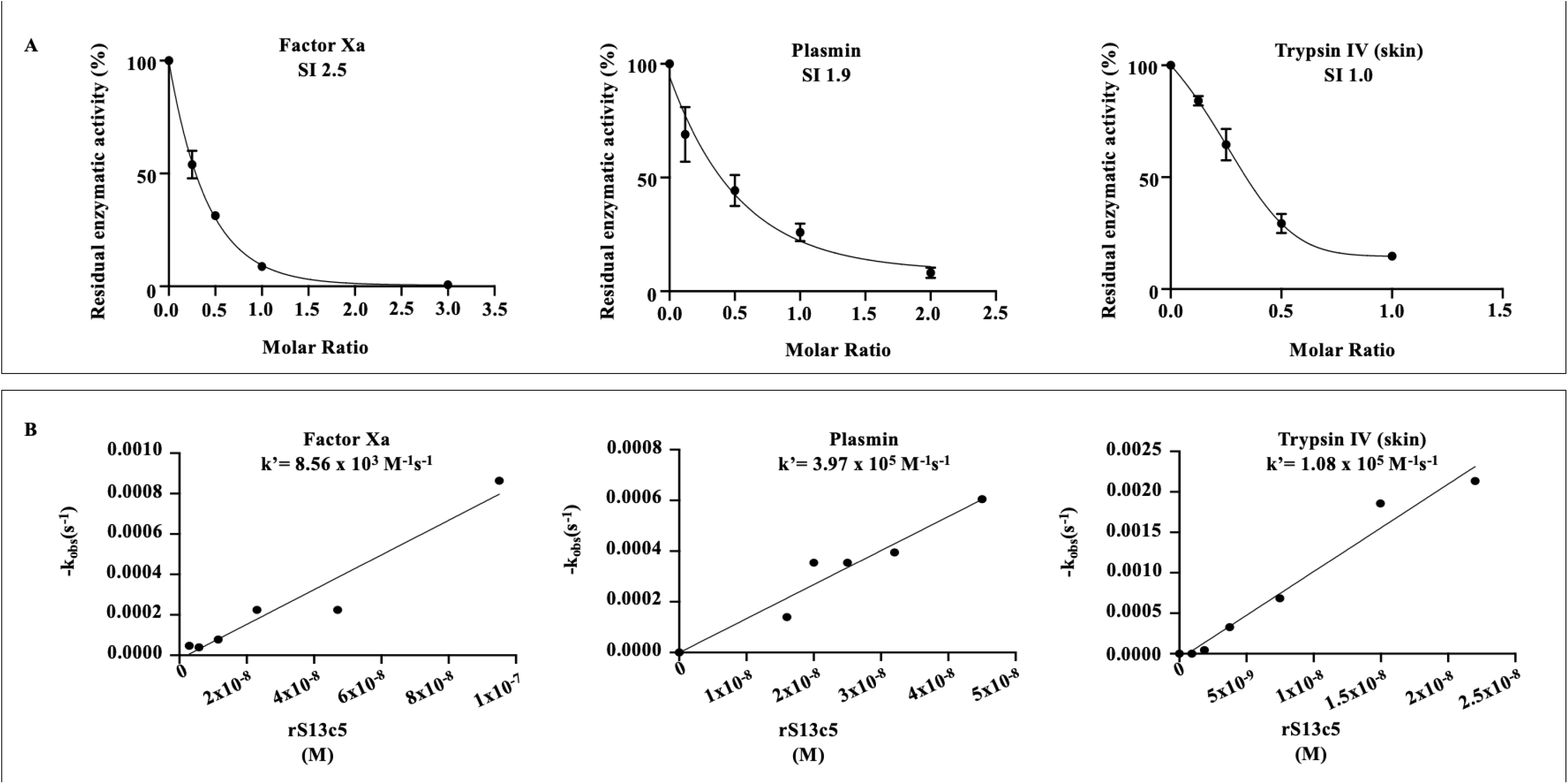
rS13c5 efficiently and rapidly inhibits plasmin and trypsin IV but exhibits slower yet effective inhibition of factor Xa (fXa). (A) Various molar ratios of rS13c5 (0-10) were incubated with fXa, plasmin, or trypsin IV for 1 h at 37 °C. Following the addition of colorimetric substrates, respective to each protease, the protease activity was recorded as outlined in materials and methods. The linear regression of the inhibitory curve was used to interpolate or extrapolate the stoichiometry of inhibition (SI: serpin-to-protease ratio) when Y = 0 as outlined. (B) Two methods were used to determine inhibition reaction rate (*K_a_*). (i) The discontinuous method was used to determine the second-order rate constant (*k_a_*) for the rate of inhibition against fXa. A constant amount of fXa (9.5 nM) was incubated with several amounts or rS13c5 (2.9-95 nM) for 0-15 min at 37 °C and residual enzymatic activity was recorded. The *k_obs_* was plotted as a function of rS13c5 concentrations. The *k_a_* for rS13c5 inhibition of fXa was determined using linear regression in PRISM 10. (ii) The continuous method was used to determine the inhibition reaction rate against plasmin and trypsin IV. A constant amount of plasmin (1.6 nM) and trypsin IV (4.74) was mixed with different amounts of rS13c5 (16.0-60 nM and 0.9-22.4 nM, respectively), and substrate (0.2 mM) was subsequently added. Substrate hydrolysis was immediately measured every 19 s for 30 min. A *k_obs_* value was calculated for each mix by nonlinear regression. The slope of the line produced from linear regression of plotted of *k_obs_* values against rS13c5 concentration provided the second-order rate constant *k*. The *k_a_* was calculated with the equation: *k_a_=k’(1+SKm)SI*.

### rS13c5 significantly delays plasma clotting time via the common pathway but not S12c5

The recalcification time assay revealed that, consistent with the substrate hydrolysis data (Fig.1), rS12c5, which did not inhibit clotting proteases, had only a modest effect on plasma clotting time (Fig. 3A), while rS13c5, which inhibited fXa, fXIa, and fXIIa, caused a dose-dependent and significant delay in clotting time (Fig. 3B). Simple linear regression analysis was successfully used to interpolate plasma clotting times in absence and presence of rS12c5 or rS13c5. For rS12c5, simple linear regression fitting of *A*_650nm_ data yielded significantly non-zero slopes (p<0.0001): Y = 0.0001271*X (control), Y = 0.0001145*X, Y = 0.0001217*X, Y = 9.853^e–005^*X, and Y = 9.846^e–005^*X for 0.28–4.5 µM rS12c5.

**Figure 3.**
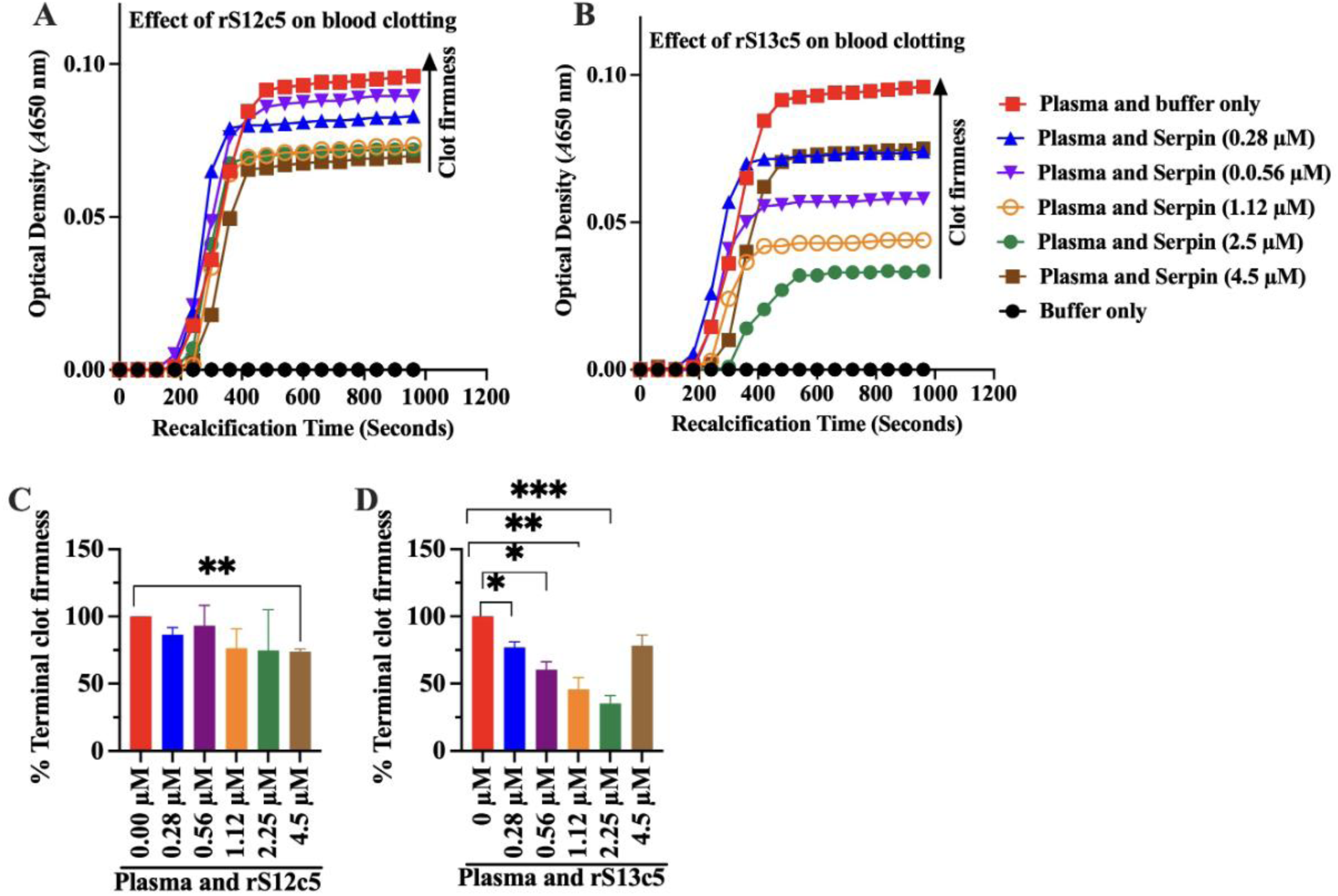
rS13c5 dose responsively delays plasma clotting and reduces the quality of the plasma clot. (A) The effects of rS12c5 and rS13c5 against blood clotting was assessed in a recalcification time assay. In triplicate reactions, 15 µL of universal coagulation reference human plasma was mixed and incubated with different amounts of rS12c5 or rS13c5 (0.28-4.5 µM) for 15 min at 37 °C. Immediately after, 150 mM CaCl_2_ was added, and clotting (recalcification) time (*A_650nm_*) was recorded every 20s for 16 min. To determine plasma clotting time (X-axis), optical density (*A*_650 nm_) data were fitted to a simple linear regression model in PRISM 10, with the Y-axis value (clotting OD) set at 0.01. The firmness or overall quality of the plasma clot (A_650nm_ data) was further analyzed using a one-phase decay model in PRISM 10 to determine the plateau. Plateau values are expressed as the percentage of clot firmness relative to the control, which is assumed to represent 100% firmness.

Assuming clot initiation at Y = 0.01, dose-dependent blood clotting start times were estimated at 78.7 s for control (or 0 µM) and 87.4, 82.2, 101.5, 101.6, and 108.8 s for various rS12c5 amounts (0.28, 0.56, 1.12, 2.25, and 4.5 µM) representing modest delays in plasma clotting of 8.7–30.1 s. Similarly, simple linear regression fitting of rS13c5 *A*_650nm_ when Y = 0.01 produced significantly non-zero linear fits (p<0.0001): Y = 0.0001271*X (control), Y = 0.0001029*X, Y = 7.967^e–005^*X, Y = 5.944^e–005^*X, Y = 4.172^e–005^\**X,* and Y = 9.684^e–005^*X for 0–4.5 µM rS13c5. Interpolated clotting start times were 78.7, 97.2, 125.5, 168.2, and 239.7 s for 0–2.25 µM, and 103.3 s at 4.5 µM corresponding to significant plasma clotting delays of 18.5–161 s and 24.6 s, respectively.

In the recalcification time assay, the increase in *A*_650nm_ optical density (OD) increased with clot firmness (56–58). In this study, one-phase decay analysis in PRISM 10 was used to determine plateau OD as the index for clot firmness (Figs. 3C, 3D). With control firmness set at 100%, rS12c5 did not significantly reduce clot quality except at the 4.5 µM dose when clot firmness was significantly reduced to 73.68% (Fig. 3C). In contrast, rS13c5 caused significant dose-dependent decline in clot firmness to 77% (p = 0.0282), 60% (p = 0.0211), 46% (p = 0.0259), and 35% (p = 0.0079) for 0.28–2.25 µM, with less effect at 4.5 µM (Fig. 3D).

### rS12c5 and rS13c5 block deposition of the MAC via the lectin pathway

The effects of rS12c5 and rS13c5 on complement activation were evaluated using the WiesLab Complement System kit (Svar Life Science #COMPL300), which enables independent assessment of the classical, alternative, and lectin pathways by measuring deposition of the membrane attack complex (MAC) (Fig. 4). Excess amounts of both rS12c5 and rS13c5 (4 µM) did not affect MAC deposition via the classical pathway (Fig. 4A), significantly reduced MAC deposition via the alternative pathway by 44% (p=0.0002) and 35% (p=0.0004) (Fig. 4B), significantly via the lectin pathway by 88% (p=0.0001) and 99% (p<0.0001) (Fig. 4C).

**Figure 4.**
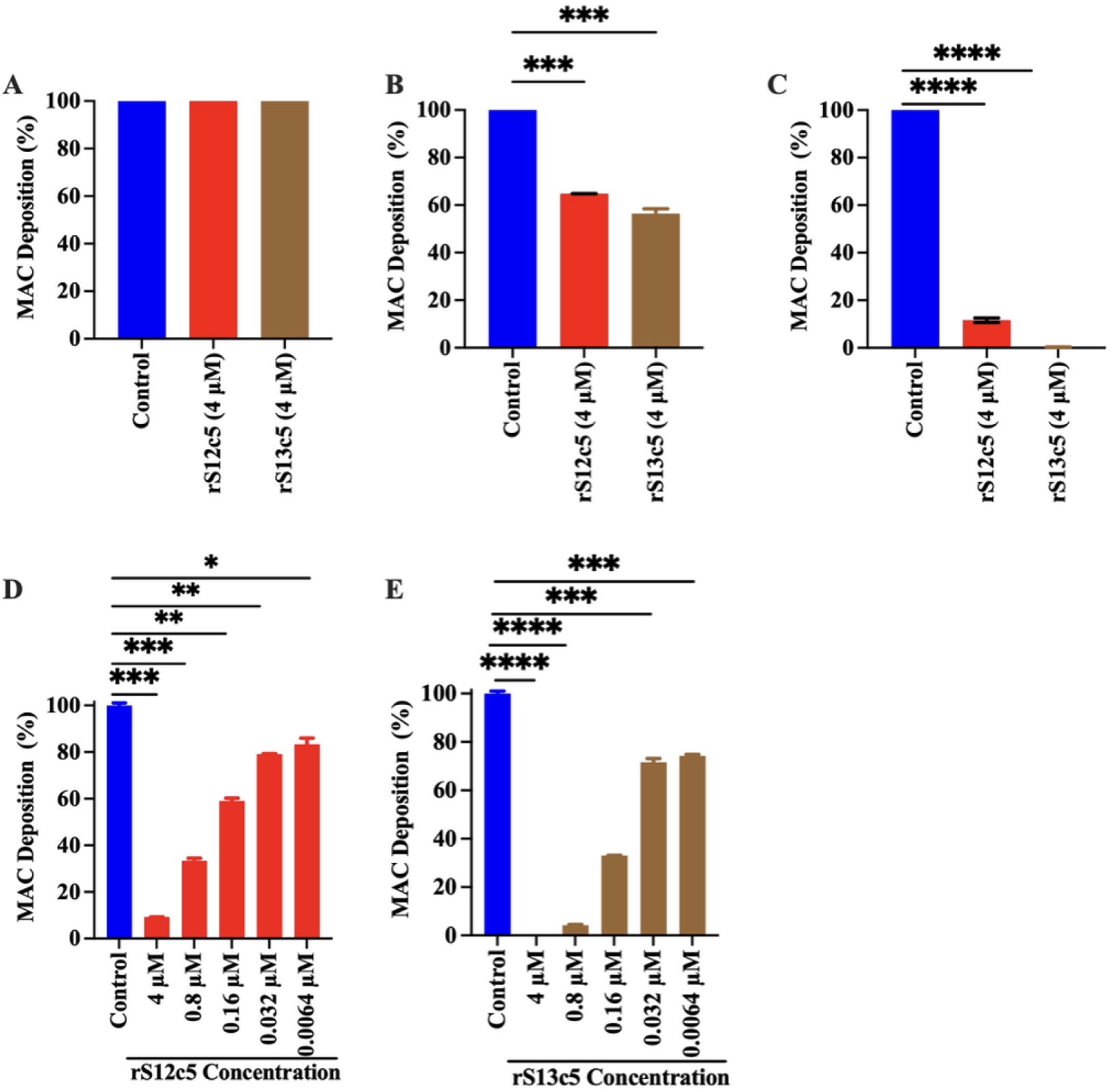
rS12c5 and rS13c5 dose-dependently inhibit complement activation via the mannose-binding lectin (MBL) pathway. (A–C) Using the WiesLab Complement System kit, excess amounts (4 µM) of rS12c5 or rS13c5 were pre-incubated with normal human serum as described under *Materials and Methods*. The effects of both proteins on activation of the (A) classical, (B) alternative, and (C) MBL pathways were independently evaluated. MAC deposition was quantified at *A*_450nm_ using a Synergy H1 plate reader. (D, E) Dose-dependent inhibition of MAC deposition via the MBL pathway was assessed using serial dilutions (4 µM – 0.0064 µM) of wild-type rS12c5 (D) and rS13c5 (E) as described under *Materials and Methods*.

We next tested the effects of serially diluted (4 to 0.0064 µM) of rS12c5 and rS13c5 in the MBL pathway (Fig. 4D and 4E). Both rS12c5 and rS13c5 inhibited MAC deposition via the MBL pathway in a dose-dependent manner. As shown, 4.0, 0.8, 0.16, 0.032, and 0.0064 µM of rS12c5 significantly inhibited MAC deposition by 90% (p<0.001), 66% (p<0.0001), 40% (p<0.0001), 20% (p=0.0001), and 16% (p=0.0005), respectively (Fig.4D). Similarly, rS13c5, 4.0, 0.8, 0.16, 0.032, and 0.0064 µM of rS12c5 and rS13c5 significantly inhibited MAC deposition by 100% (p<0.001), 95% (p<0.0001), 66% (p<0.0001), 28% (p<0.0001), and 25% (p<0.0001), respectively (Fig. 4E).

### rS12c5 and rS13c5 inhibit complement activation by binding MASP1/3 and other complement factors

We next sought to determine the mechanisms by which S12c5 and S13c5 inhibit membrane MAC deposition. We performed pull-down assays and subjected pull down eluates to western blotting analysis (Figs. 5A-5H) and ELISA analysis (Figs.5I-5O) using antibodies to specific complement proteins. For western blotting analysis assays, the antibody to the Histidine recombinant protein fusion tag was used to validate S12c5 (lanes 3 and 4) and S13c5 (lanes 1 and 2) in reactions (Fig.5A). Both rS12c5 and rS13c5 interacted with the C1q and C1s components of the complement C1 complex (Fig. 5B and 5C, lanes 2 and 4), whereas C1r interacted only with rS13c5 (Fig. 5B, lane 2). Similarly, complement proteins, C3, C5, C9, and MASP-1/3 bound to both rS12c5 and rS13c5 (Figs 5D-5H, lanes 2 and 4). Notably, with the exception of the antibody to C1s, other antibodies to complement proteins did not cross-react with rS12c5 and rS13c5 (Fig.5B-5H, lanes 1 and 3).

**Figure 5.**
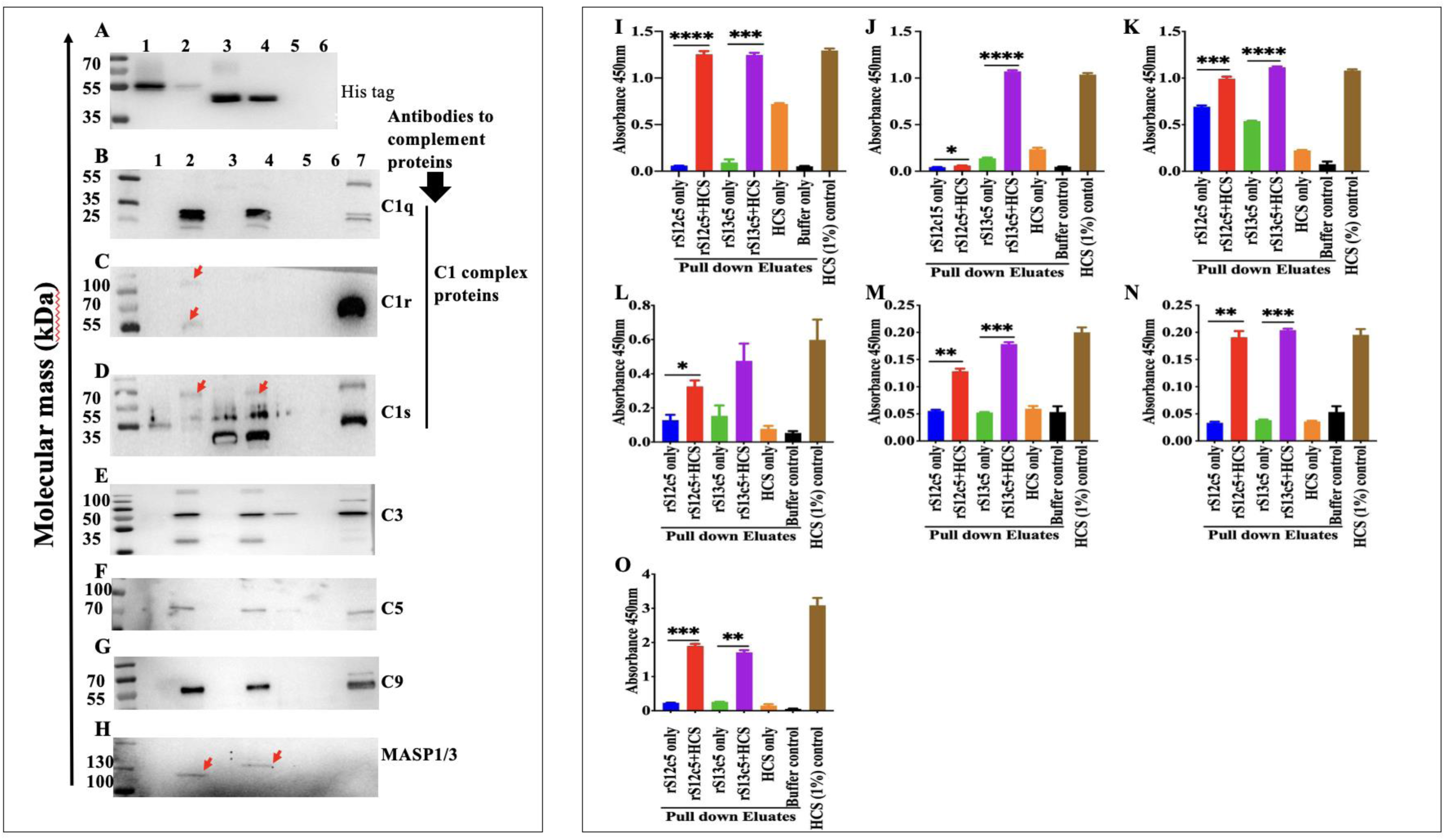
rS12c5 and rS13c5 inhibit complement activation through interactions with Mannan-binding lectin-associated serine proteases (MASP)-1/3 and other complement components, as revealed by pull-down assays. To identify human complement proteins that interact with rS12c5 and rS13c5, Dynabeads™ His-Tag Pulldown magnetic beads were coated with 10 µg of affinity-purified rS12c5 or rS13c5. The coated beads were then incubated with 1% human complement serum (HCS). Western blotting analysis of eluates using the antibody to the Histidine recombinant protein fusion tag (Fig. 5A) confirmed rS13c5 (lanes 1 and 2) and rS12c5 (lanes 2 and 4). Eluates were subjected to western blotting (Figs.5B-5H) and ELISA analysis (Figs.5I-5O) using antibodies to complement proteins. Western blot lanes 1–7 respectively represent: rS13c5 only (1), rS13c5 + HCS pull-down eluate (2), rS12c5 only (3), rS12c5 + HCS pull-down eluate (4), HCS only control (5), buffer only control (6), and positive control (1% HCS [7]). Figures 5I–O show ELISA analyses of pull- down eluates probed using antibodies to C1q, C1r, C1s, C3, C5, C9, and MASP-1/3, respectively.

Notably, apparent complexes between the activated C1r B chain and rS13c5 were detected (red arrowhead, Fig. 5C lane 2), but not for rS12c5 (Fig. 5C lane 4). Conversely, the activated C1s B chain apparently formed complexes with both serpins (red arrowheads, Fig 5D lanes 2 and 4). The C1r antibody detected two bands at approximately 55 kDa and 120 kDa in the rS13c5 pull-down eluate (Fig.5C, lane 2, red arrowhead). The upper band (∼120 kDa) corresponded to the expected molecular weight of a complex between glycosylated rS13c5 (70–100 kDa; Supplementary Fig. S2C) and the 27 kDa serine protease B chain of activated C1r (59). Similarly, western blots probed with antibody to C1s showed a distinct ∼70 kDa band, in addition to some nonspecific binding to both serpins (Fig.5D, lanes 2 and 4). The ∼70 kDa band is consistent with a complex between the 28 kDa activated C1s serine protease B chain (60) and the less glycosylated forms of the serpins (∼50 kDa; Supplementary Figs. S2A, S2B). Furthermore, the antibody to MASP-1/3 detected high–molecular-weight bands at approximately 120 kDa and 130 kDa for rS13c5 and rS12c5, respectively, consistent with complexes between either serpin and MASP-1 or MASP-3 (∼90 kDa; (61, 62) (Fig. 5H lanes 2 and 4, red arrowhead).

ELISA analyses of pull-down eluates supported the western blot results, showing that both S12c5 and S13c5 interacted with complement serine proteases (C1r, C1s, MASP-1/3) (Figs. 5J, 5K, and 5O) as well as non-protease components (C1q, C3, C5, and C9) (Figs 5I, 5L-5N). Consistent with western blotting data, ELISA confirmed strong binding of C1r to rS13c5 but not rS12c5 (Fig. 5J).

### LC–MS/MS analysis of pull-down eluates provides additional functional insights into rS12c5 and rS13c5

To gain further functional insights into S12c5 and S13c5, pull-down eluates were analyzed by LC-MS/MS (Supplementary Table S1). Tandem mass spectra (MS/MS) were searched against a human protein database from GenBank. Proteins supported by at least two distinct MS/MS spectra (or inferred peptides) were considered high-confidence and prioritized for analysis. From the rS12c5 pull-down eluate, 356 proteins were identified, including 177 absent in the control (serum only) eluate. Similarly, 471 proteins were detected in the rS13c5 pull-down eluate, with 189 uniquely present in this sample (Supplementary Table S1). These datasets were subjected to pathway enrichment analysis using the Reactome platform (63). Significant enrichment (p < 0.05) was observed for 27 pathways in the rS12c5 dataset and 47 pathways in the rS13c5 dataset (Supplementary Tables S2 and S3). In both cases, proteins were strongly associated with complement activation and other innate immune system pathways. Specifically, rS12c5 proteins significantly mapped to five complement-related pathways: Terminal pathway of complement (R-HSA-166665), Regulation of complement cascade (R-HSA-977606), Complement cascade (R-HSA-166658), Classical antibody-mediated complement activation (R-HSA-173623), and initial triggering of complement (R-HSA-166663). Similarly, rS13c5 proteins mapped to seven complement-related pathways, including the same five as for rS12c5, plus Creation of C4 and C2 activators (R-HSA-166786) and Activation of C3 and C5 (R-HSA-174577). The relative abundance of complement system factors identified in both eluates is summarized (Fig. 6). Notably, the C1 protease inhibitor (a serpin regulator of complement activation) was detected in both rS12c5 and rS13c5 eluates. It is also notable that most C1s, C6, C7, and C8 (alpha, beta, and gamma) were exclusively detected in rS12c5 and S13c5 eluates and not in controls (Fig.6). Interestingly, although western blotting and ELISA confirmed the presence of MASP1/3 in both S12c5 and S13c5 eluates, these proteins were not detected in the LC-MS/MS datasets and the high stringency screening needed a two-peptide hit. However, in the preliminary screen, MASP1/3 was detected at low stringency with a one peptide hit.

**Figure 6.**
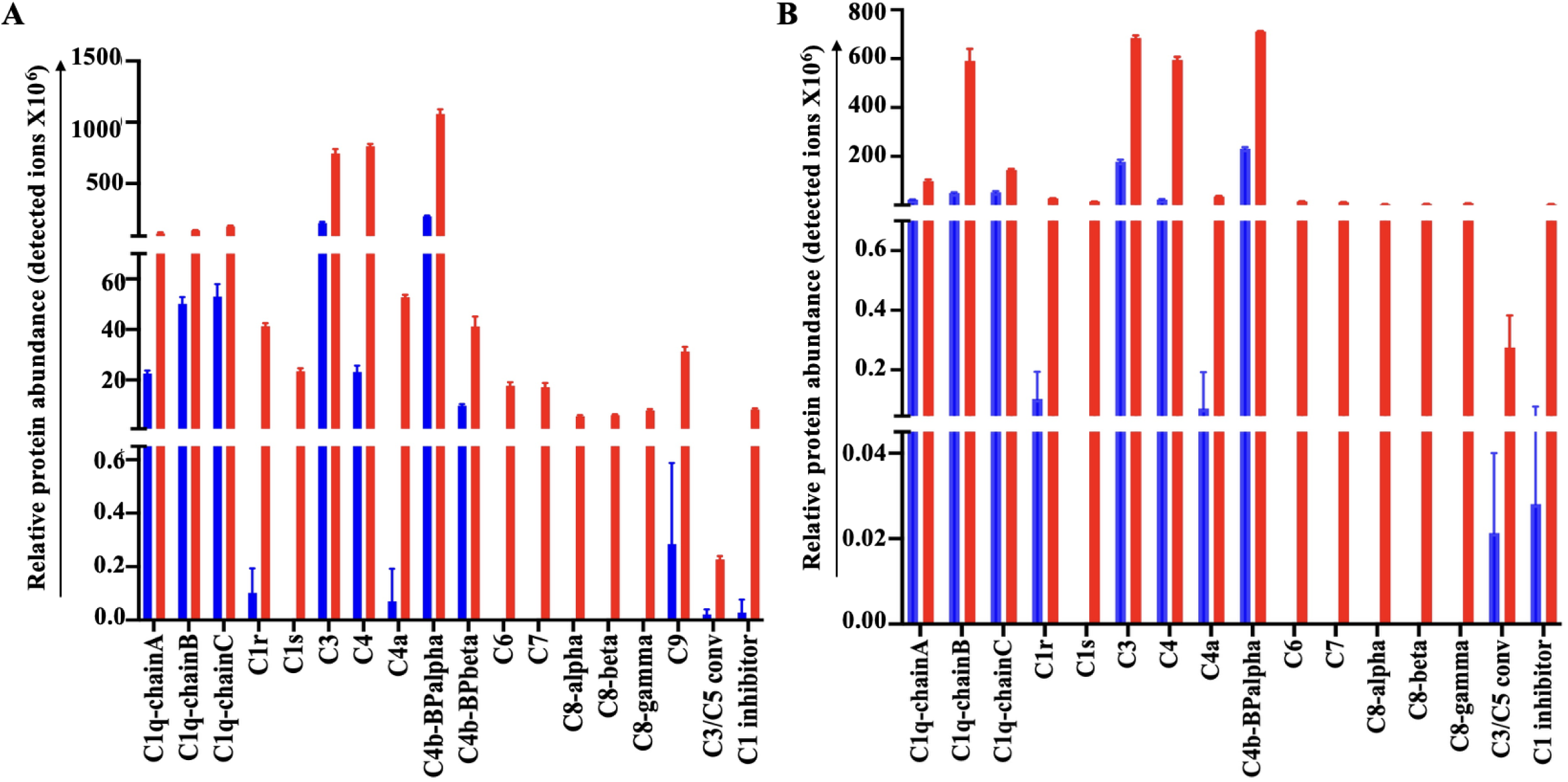
LC-MS/MS analysis reveals complement system proteins interacting with rS12c5 and rS13c5. Pull-down eluates were subjected to LC-MS/MS analysis to identify human complement proteins interacting with rS12c5 and rS13c5. Samples were prepared using the in-solution digestion approach as described in the *Materials and Methods* section. MS/MS analysis was performed using an Orbitrap Fusion Tribrid Mass Spectrometer (Thermo Fisher Scientific, Bremen, Germany) coupled to a Dionex UltiMate 3000 reverse-phase nano-UHPLC system (Thermo Fisher Scientific, Bremen, Germany). The resulting tandem mass spectra were searched against the human protein database downloaded from GenBank to identify proteins present in the pull-down eluates (Supplemental Table 1). Complement system proteins identified as interactors of rS12c5 (A) and rS13c5 (B) are summarized. The Y-axis represents the relative abundance of complement proteins (red bars) compared with those detected in human complement serum (HCS) only controls (blue bars), based on protein ion counts.

While rS12c5 had only a modest effect on blood clotting, its pull-down eluate contained several coagulation system proteases, including thrombin and factors IX (fIX), XI (fXI), and XII (fXII), a profile like that of rS13c5, which significantly interfered with clot formation through the common pathway (Supplementary Table S1). Notably, although substrate hydrolysis assays showed that rS13c5 strongly inhibited factor Xa (fXa), this protein was not detected under high-stringency conditions requiring two peptide matches; however, it was identified under low-stringency conditions requiring one peptide match in the rS13c5 eluate (Supplementary Table S1).

Another notable feature of the LC-MS/MS data was the concurrent pull down of proteases and their corresponding regulatory inhibitors (Supplementary Table S1). Both rS12c5 and rS13c5 eluates contained complement system proteases (C1r, C1s, and MASP1/3) along with the C1 inhibitor, the regulatory serpin of the complement system along with contact coagulation, and fibrinolysis (64). Additionally, inter-alpha-trypsin inhibitor, a multifunctional non-serpin inhibitor that also regulates complement activation among other functions (65) was detected in both pull down eluates. Similarly, both proteins pulled down thrombin along with its inhibitor, antithrombin (66), and kallikrein together with its regulatory serpin, kallistatin (67). Also detected in both eluates was serpin A5 also known as protein C inhibitor, a multifunctional serpin that regulates several serine proteases such as activated protein C and coagulation factors (fII) thrombin, fXa, and fXIa (67). The mass spectrometry proteomics data have been deposited to the ProteomeXchange Consortium (Dataset Identifier: PXD074437) .

### Molecular modeling predicts that S13c5 interacts with complement proteases more efficiently than S12c5

Recently, a co-crystal structure of the endogenous SERPIN regulator of the classical and lectin pathways (C1 esterase inhibitor, C1-INH), in complex with complement C1s has been reported (PDB:8W18, (68). The C1s/C1-INH structure involves the initial noncovalent interaction formed prior to SERPIN activation (*i.e.*, encounter complex/ “Michaelis-Menten complex”). To better understand potential molecular mechanisms of tick serpin-mediated complement inhibitory activity, we used AlphaFold3 to model the putative encounter complexes formed by tick serpin proteins (S12c5 and S13c5) and serine proteases of the complement cascade.

We began by modeling S12c5 and S13c5 with the catalytic serine protease (SP) domain of human C1s (Figs. 7A and 7B). AlphaFold3 produced a high-quality fold of each individual protein component as judged by pLDDT scores > 90 for a majority of residues in S12c5 and S13c5, and C1s. S12c5 and S13c5 adopt canonical SERPIN folds as evidenced by low root mean square deviation (RMSD) values when aligned to C1-INH in PDB: 8W18 (S19 RMSD = 1.78 Å; S20 RMSD = 1.58 Å). The SP domain of C1s in each model also agrees closely with the experimentally determined structure, with RMSD values of 0.442 Å and 0.433 Å in the C1s/S13c5 and C1s/ S12c5 models, respectively. For C1s/S13c5, the complex is also confidently predicted, as judged by an interface predicted modeling (iPTM) score > 0.8, and generally low values in the plot for the predicted aligned error (PAE) (Appendix Fig. S3A1-8). Furthermore, the C1s/S13c5 interface predicted by the model is consistent with the expected engagement of the S13c5 RCL with the active site of C1s as observed in C1s/C1-INH and other experimentally determined SERPIN/protease encounter complexes (68). In contrast, the S12c5/C1s model is poorly predicted by all measures including exhibiting a very low iPTM score (0.19) and a non-canonical SERPIN/protease complex (Fig. S3B1-8).

**Figure 7.**
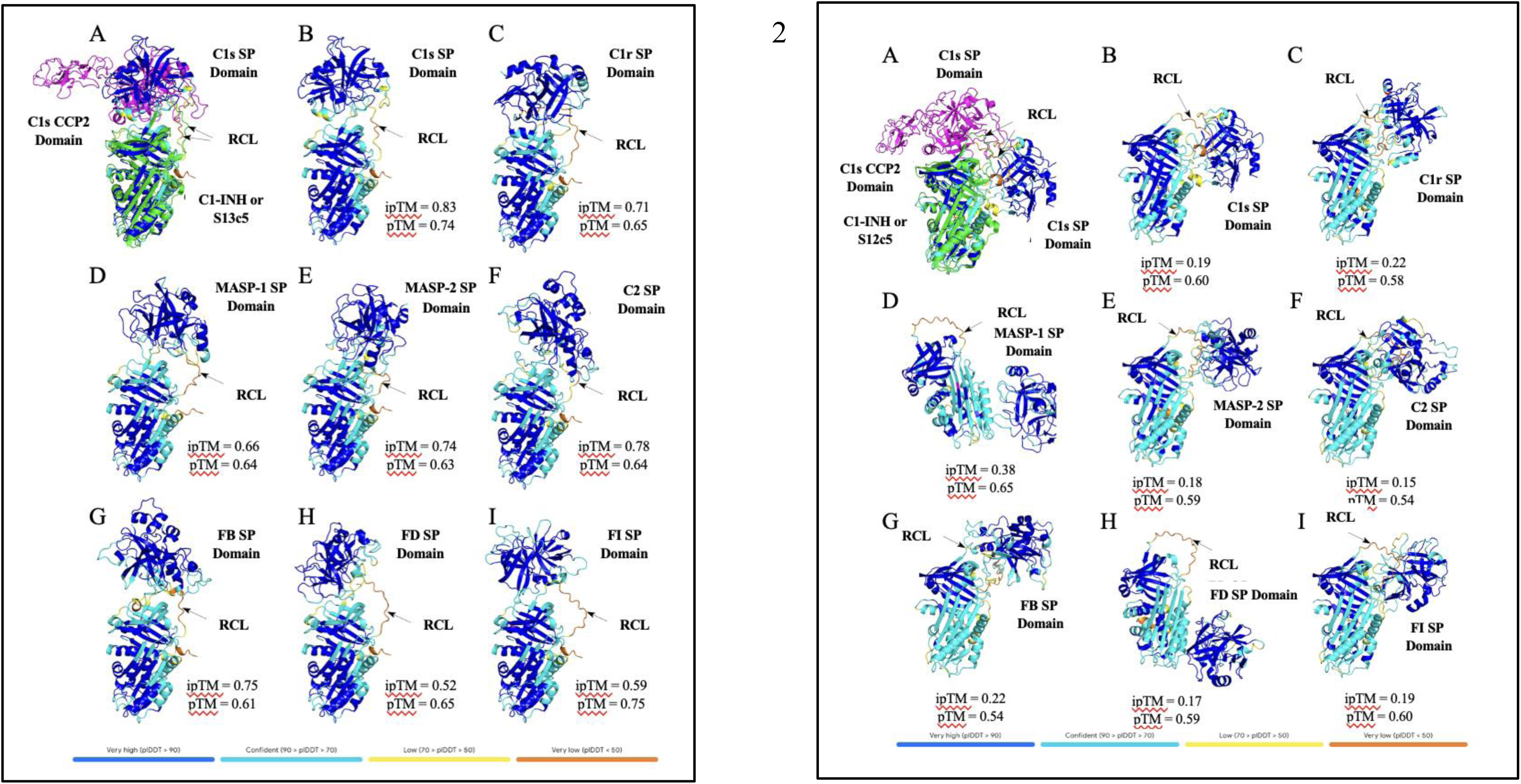
AlphaFold3 modeling predicts high confidence interactions between complement serine proteases and S13c5 (7 panel 1), but not with S12c5 (7 panel 2). In Fig. 7 panel 1, (A-I) AlphaFold3 models resulting from co-folding the S13c5 protein sequence with sequences corresponding to the serine protease (SP) domains of the complement proteases are shown. Panel A depicts a structural alignment between the AlphaFold3-predicted C1s/S19 model (panel B) and the published co-crystal structure of C1s in complex with the native human SERPIN regulator, C1-INH (PDB:8W18) For the C1s/C1-INH crystal structure C1s is colored magenta and C1-INH in green. Note that in PBD:8W18 the C1s construct included the complement control protein 2 domain (CCP2). The reactive center loop (RCL) of C1-INH or S13c5 is indicated by an arrow. Per-residue confidence scores, reported as predicted Local Distance Difference Test (pLDDT) values, are displayed on the model using AlphaFold3’s default color scheme. pTM: Predicted template modeling score. ipTM: interface predicted template modeling score. In Fig. panel 2, AlphaFold3 models resulting from co-folding the S12c5 protein sequence with sequences corresponding to the serine protease (SP) domains of the complement proteases are shown. The layout is identical to that shown in Fig. 7 panel 2.

This approach was carried out using seven additional serine proteases of the complement system: C1r, mannan-binding lectin-associated serine protease 1 (MASP-1), mannan-binding lectin-associated serine protease 2 (MASP-2), C2, Factor B (FB), Factor D (FD), and Factor I (FI) (Fig. 7A C-I). Based on iPTM scores alone, S13c5 co-folds produce moderate-quality models (*i.e.*, iPTM > 0.6 and < 0.8) with C1r, MASP-1, MASP-2, C2, and FB. Each of these complexes were also generally consistent with a canonical serpin/protease encounter complex (Fig. 7A, C-G). Conversely, the models of FD/S13c5 and FI/S13c5 produced poor quality models (iPTM < 0.6) (Fig. 7A, H-I). Interestingly, and in stark contrast to S13c5, all models of complement protease complexes with the S12c5 SERPIN produced low quality predictions, including C1s (Figs. 7B and Appendix Fig. S3B)

### rS12c5 and rS13c5 protect complement-sensitive B. burgdorferi from complement mediated killing

Prompted by our anti-complement function results, we next tested whether rS12c5 and rS13c5 could protect the complement-sensitive *B. burgdorferi* strain B314 using a serum sensitivity assay (Fig. 8). As a positive and negative control, we incubated recombinant *B. miyamotoi* FbpA (BmA) protein and mutant form of BmA that, in stark contrast to intact BmA, cannot bind or inhibit human C1r—designated BmA DA (69)—in normal human serum (NHS) prior to incubation with serum *B. burgdorferi* strain B314. This was compared relative to untreated NHS. While the untreated and the NHS incubated with BmA DA resulted in killing of *B. burgdorferi* strain B314, the addition of BmA provided significant protection to this serum sensitive borrelial strain (Fig. 8). In addition, both intact forms of rS12c5 and rS13c5 significantly rescued B314 from complement-mediated killing. At a concentration of 4 µM, rS12c5 and rS13c5 increased B314 survival by more than 80% and 60%, respectively, with both effects reaching high statistical significance (p < 0.0001). Importantly, this protective activity was abolished when using either heat-inactivated proteins or RCL deletion mutants, confirming that the observed rescue was specific to the intact, functional serpins.

**Figure 8.**
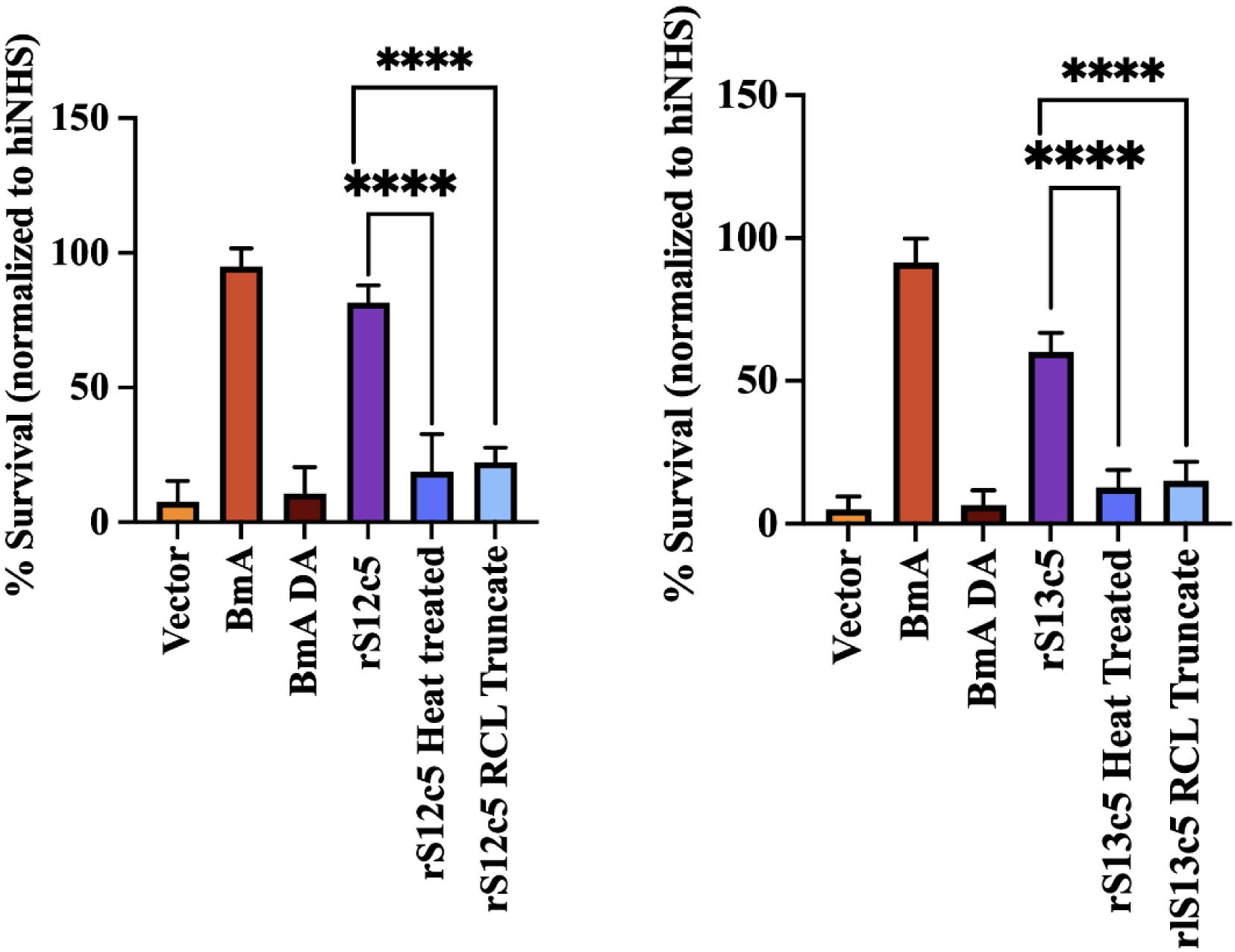
Treating serum with rS12c5 and rS13c5 rescues sensitive *Bb* from complement. Normal human serum was pre-incubated without (no treatment control, far left sample shown for both plots) or with added wildtype *B. miyamotoi* FbpA (BmA; at 0.25 mM) that binds and inhibits human C1r, a mutated form of BmA (designated as BmA DA) that does not bind or inhibit human C1r, S12c5 (left panel) or S13c5 (right panel), a heat inactivated (HI) of rS12c5 or rS13c5 (4 µM; treated for 30 min at 37 °C), and a reactive center loop (RCL) deletion mutants of both S12c5 and S13c5. After preincubation with the different proteins in normal human serum, the serum samples were added to serum-complement sensitive *B. burgdorferi* strain B314 at concentration of 1x10^6^ spirochetes in their respective wells and incubated at 32 °C at 100 rpm for 3 h. These assays were done in triplicate. Following incubation, Bb survival was scored by counting spirochetes in 10 fields of view under dark field microscopy. Data are presented as mean % survival normalized to hiNHS ± SEM.

### rS13c5 enhances B. burgdorferi colonization of C3H/HeN mouse organs

Given the broad functional activity of rS13c5, we next examined its role in Bb dissemination and tissue colonization in C3H/HeN mice. Female mice were intradermally co-inoculated (CI) with rS13c5 (25 µg) and the Bb strain B31 MSK5 derivative (70, 71) at either 1 × 10⁴ or 1 × 10⁶ spirochetes (Fig. 9). At 7 days post-CI (PCI), mice co-inoculated with rS13c5 and 1 × 10⁶ spirochetes showed a 2.3-fold increase in spirochete burden in ear skin compared to controls, though this difference was not statistically significant. By 14 days PCI, co-inoculation with rS13c5 and 1 × 10⁴ spirochetes resulted in a significantly higher bacterial load in ear skin (13.5-fold increase, *p* < 0.0001) and joint tissue (3.4-fold increase, *p* = 0.0367). Additional increases were observed in inguinal lymph nodes (2.0-fold), bladder (4.9-fold), and dura mater (2.1-fold), although these were not statistically significant. In contrast, no significant differences were detected in bladder, heart, or dura mater when rS13c5 was co-inoculated with 1 × 10⁶ spirochetes at 14 days PCI.

**Figure 9.**
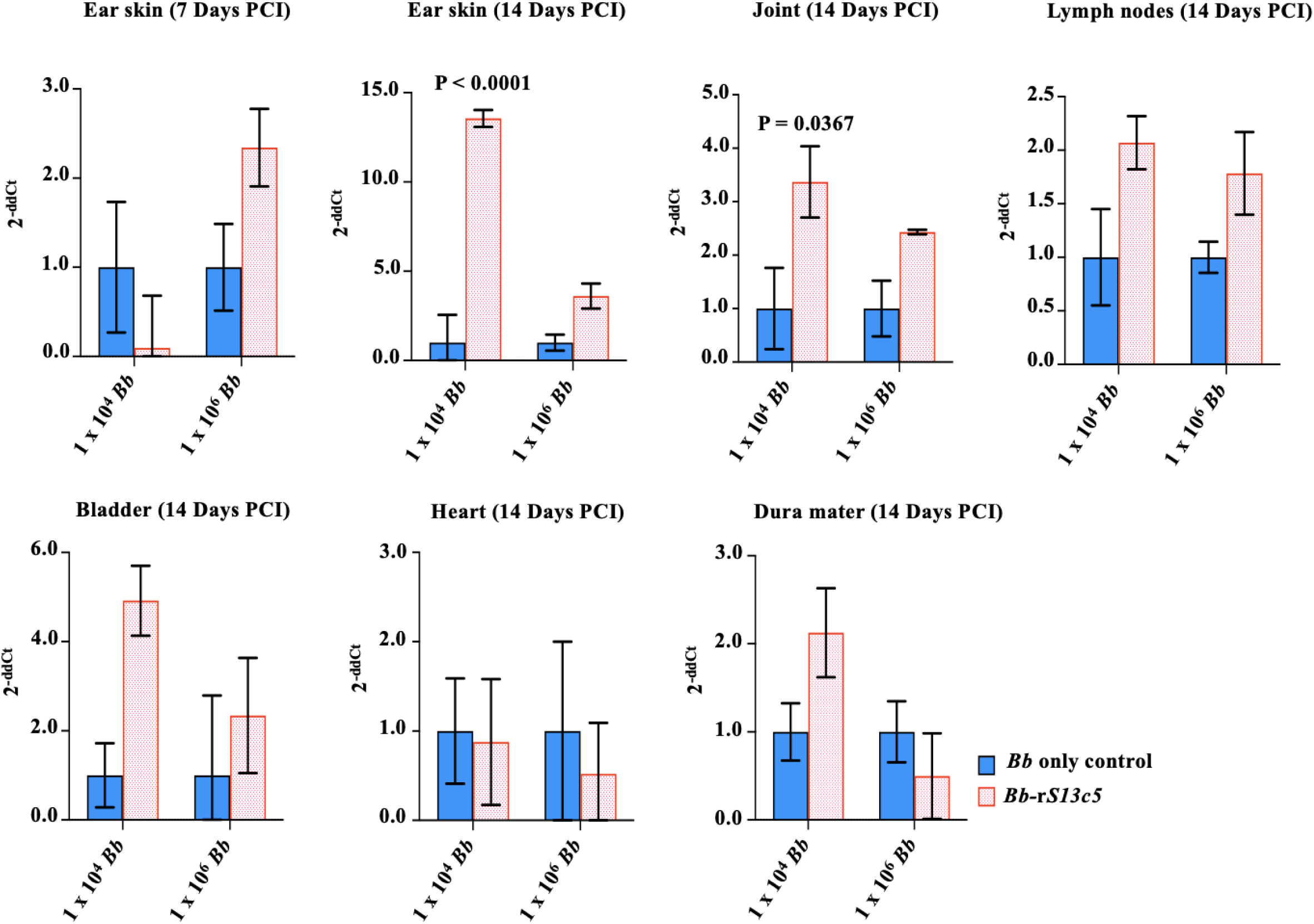
Co-inoculating rS13c5 and *Bb* strain 31 enhanced spirochete colonization of mice ear skin and joint. Three 9–10-week-old female C3H/HeN mice per treatment were needle inoculated (intradermally) with 1x10^4^ *Bb*, 25 µg of rS13c5 and 1x10^4^ *Bb*, 1x10^6^ *Bb*, or 25 µg of rS13c5 and 1x10^6^ *Bb.* Punch biopsies of ear skin were taken 7 days PCI. Mice were euthanized by CO_2_ 14 days post co-inoculation (PCI) and the following organs were obtained for gDNA extraction: ear skin, joint, inguinal lymph nodes, bladder, heart, and dura mater. Approximately 5-10 ng of gDNA were used for qRT-PCR quantification of *Bb* using *Bb flaB* and murine *β*-actin (internal reference gene) primers. The data are presented as fold change (2^-ΔΔCt^) of *Bb flaB* gene amongst treatments compared to the *Bb* only control group.

Because Bb sensu stricto is known to disseminate to joints and cause Lyme arthritis (72), we also assessed joint swelling between the tarsocrural joint and calcaneus bone (Fig. 10A). At 7 days PCI, mice inoculated with 1 × 10⁴ spirochetes alone showed greater swelling (30.42% ± 13.07, not significant [ns]) compared to mice co-inoculated with rS13c5 and 1 × 10⁴ spirochetes (9.94% ± 3.70, ns). Similarly, at 14 days PCI, mice given 1 × 10⁶ spirochetes alone displayed greater swelling (27.98% ± 9.62, ns) than those co-inoculated with rS13c5 and 1 × 10⁶ spirochetes (21.83% ± 4.50, ns) (Fig. 10B). Consistent with these findings, joint swelling at 14 days PCI was higher in mice inoculated with 1 × 10⁴ (30.0% ± 15.28, ns) or 1 × 10⁶ spirochetes (35.98% ± 12.34, ns) compared to those co-inoculated with 25 µg of rS13c5 and either 1 × 10⁴ (17.9% ± 2.50, ns) or 1 × 10⁶ spirochetes (15.4% ± 7.70, ns), respectively (Fig. 10B).

**Figure 10.**
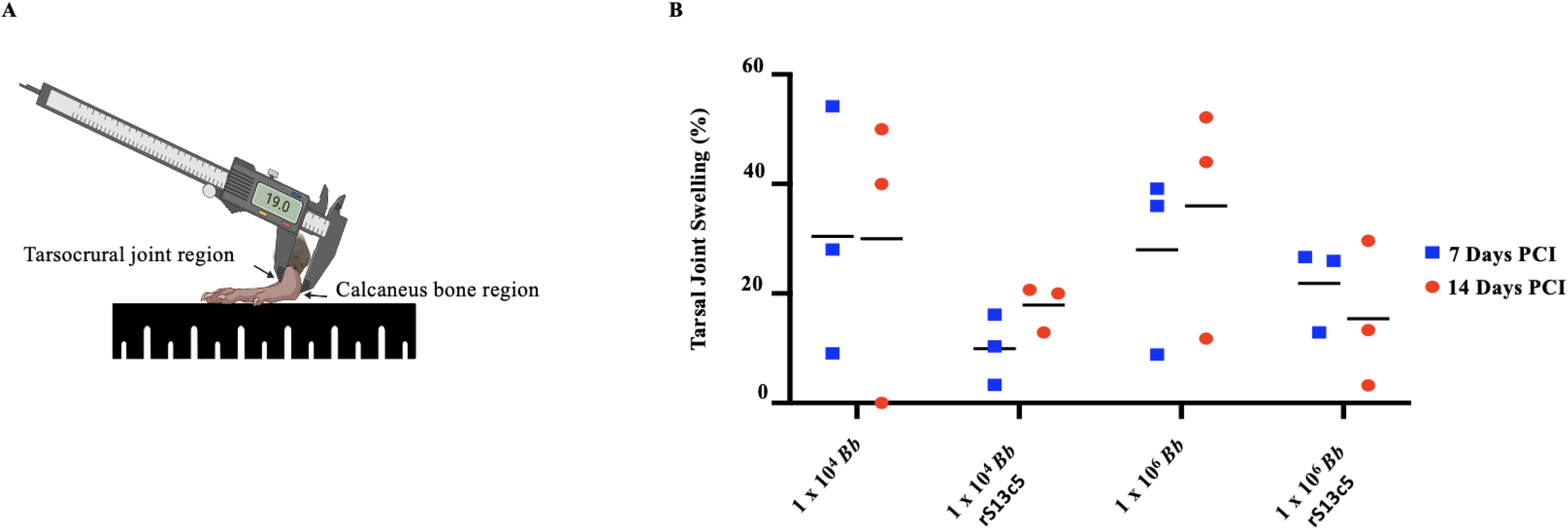
Co-inoculation of rS13c5 and Bb strain 31 decreased tarsal joint swelling. (A) Both tarsal joints were measured pre-treatment (normal tarsal joint), seven-, and fourteen-days post co-inoculation. (B) The data are presented as mean of tarsal joint swelling percent ± SEM.

### S12c5 is highly immunogenic compared to S13c5

Because S12c5 and S13c5 are secreted into the host during feeding by uninfected and Bb*-*infected *I. scapularis* nymphs (24), we investigated their immunogenicity using immune sera (for IgM testing) and purified IgG from naïve rabbits (pre-immune) and from rabbits fed on by either uninfected or Bb–infected *I. scapularis* nymphs (Figs. 11 and 12). Western blot analysis showed modest IgM binding to glycosylated (DG–) rS12c5 and rS13c5 in sera from both uninfected tick- (Fig. 11A2 and 11B2) and Bb-infected tick-(Figs. 11A3 and 11B3) fed rabbits, whereas no binding was detected to deglycosylated (DG+) proteins (Figs. 11C1–D3). In contrast, purified IgG displayed differential binding patterns. IgG bound strongly to both glycosylated rS12c5 (Figs. 11E2, E3) and weakly to glycosylated rS13c5 (Figs. 11F2, F3). Similar binding patterns were observed for deglycosylated proteins (Fig. 11G2 and G3, H2 and H3). Notably, deglycosylation markedly reduced IgG binding intensity to both proteins (Fig. 11G–H). Furthermore, IgG from rabbits fed on by uninfected ticks showed stronger binding to deglycosylated proteins (Fig. 11G2, 3H2) compared with IgG from infected tick-fed rabbits (Fig. 11G3, 3H3). As expected, neither IgM nor IgG from naïve rabbits bound to S12c5 or S13c5, confirming assay specificity.

**Figure 11.**
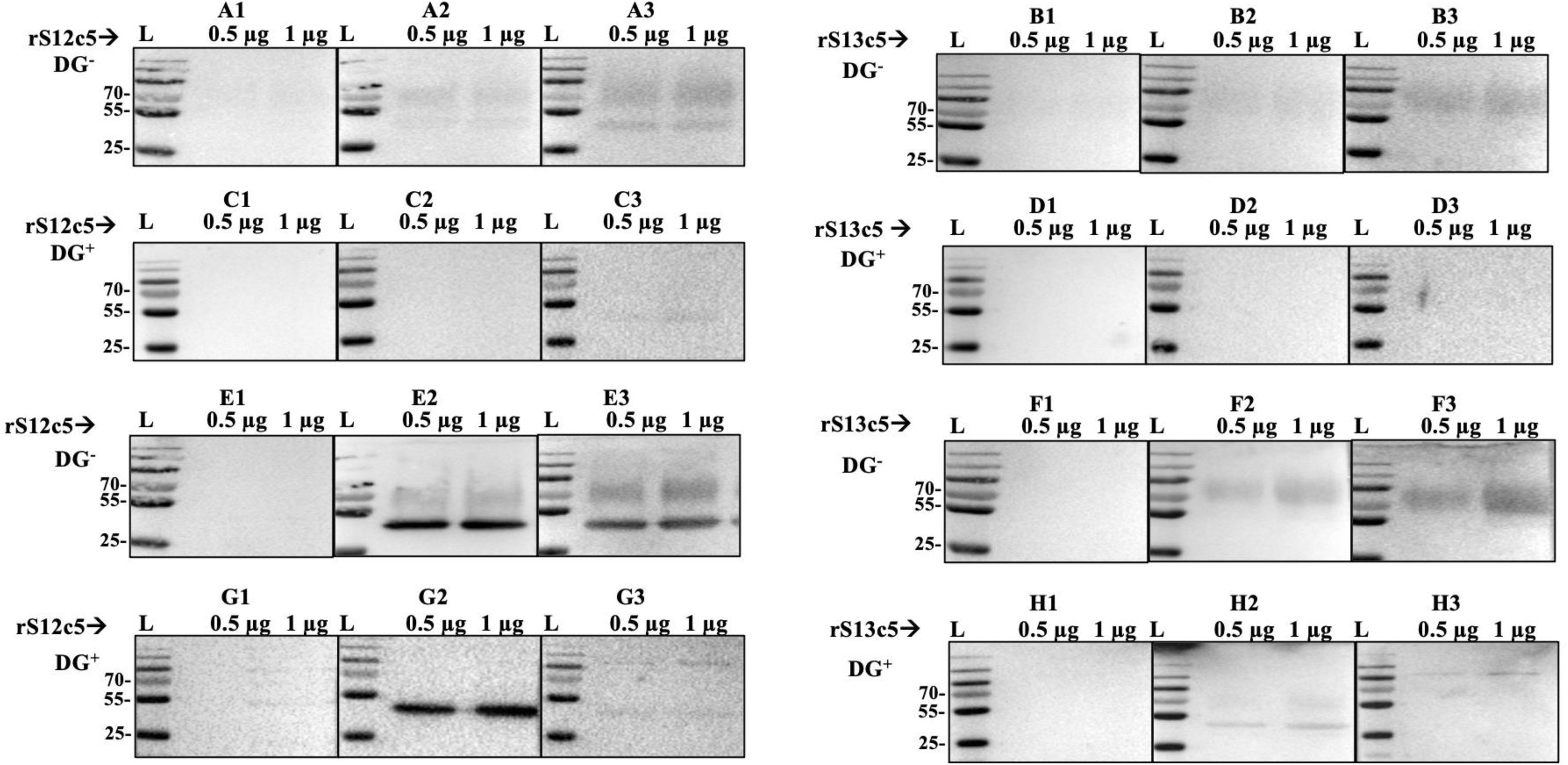
S12c5 and S13c5 are immunogenic. Affinity purified glycosylated (DG^-^) or deglycosylated (DG^+^) rS12c5 and rS13c5 (0.5 and 1 µg) were subjected to standard western blotting analysis using the indicated antibodies as detailed under Materials and Methods. For IgM binding, membranes were probed with 1:100-diluted rabbit serum collected at one week after uninfected or Bb**-** infected *I. scapularis* nymph tick feeding (A-D). For IgG binding, membranes were probed with empirically optimized 16 µg of purified IgG from sera collected at two weeks after uninfected or Bb***-***infected feeding (E-H). After appropriate washing and exposure to HRP conjugated IgM or IgG secondary antibodies (1:5000 diluted), antibody binding was detected with chemiluminescent substrates and visualized using the ChemiDocXRS+ imager (Biorad).

**Figure 12.**
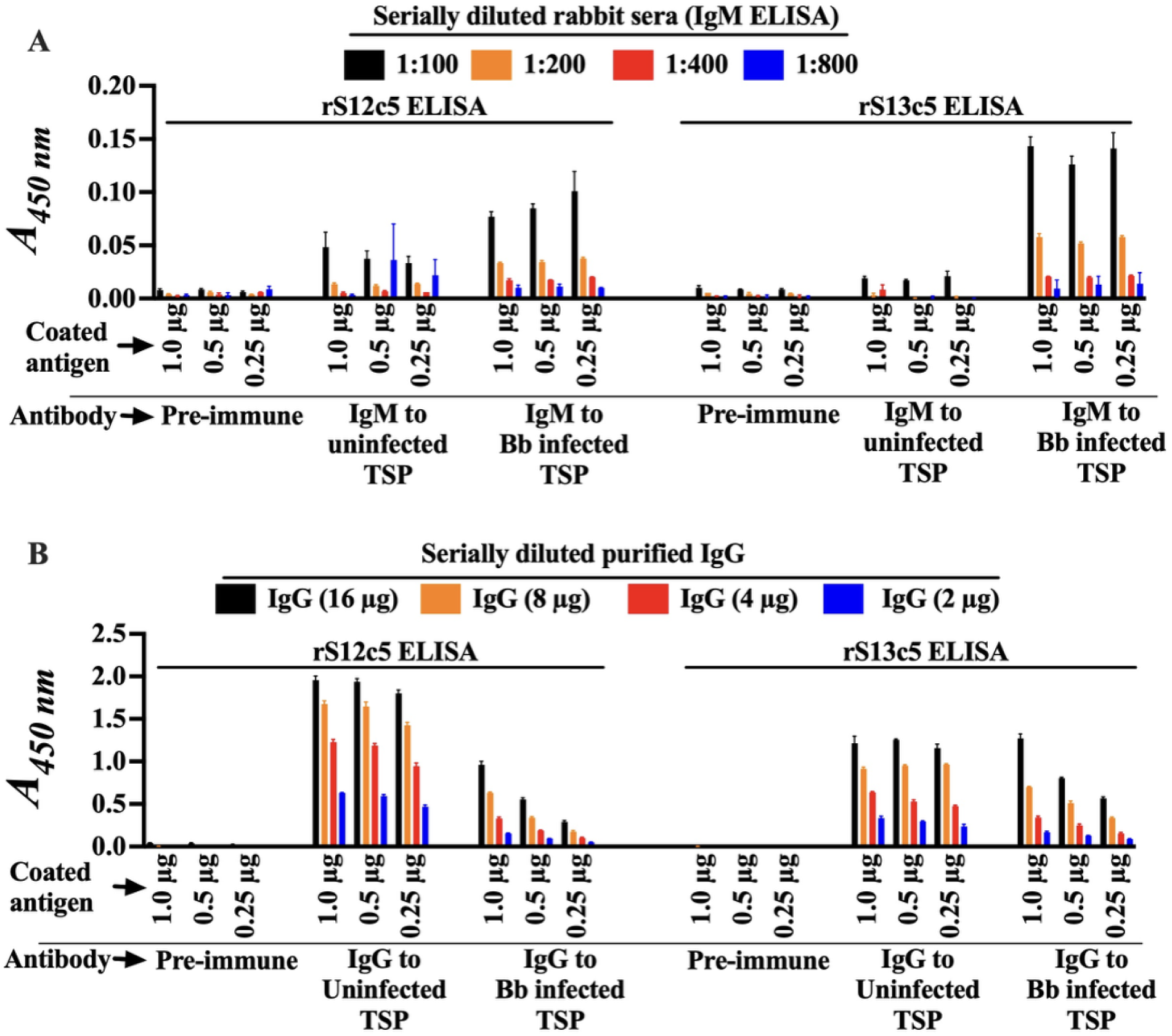
Antibody responses in rabbits fed on by uninfected and Bb–infected ticks show higher titers to the less active rS12c5 than to the more active rS13c5. Various concentrations of rS12c5 and rS13c5 (1.0–0.25 µg) were subjected to standard ELISA using serial dilutions of rabbit immune sera (1:100–1:800 for IgM detection) or purified IgG (16–2 µg), as described in the *materials and methods*. After washing, plates were incubated with goat anti-rabbit IgM or IgG (1:2000), followed by substrate addition for 15 min. The reaction was stopped with sulfuric acid, and absorbance was measured at *A*_405nm_ using a Synergy H1 plate reader. Data are presented as mean A405 ± SEM.

ELISA analysis further supported these findings (Fig. 12). S12c5 was more immunogenic than S13c5, as reflected by higher absorbance values. IgM titers to both proteins were significantly elevated in rabbits fed on by Bb-infected ticks compared to those fed on by uninfected ticks (Fig. 12A), consistent with protein secretion dynamics (24). In contrast, IgG responses did not parallel these dynamics (Fig. 12B). Rabbits fed on by uninfected ticks developed higher IgG titers to rS12c5 than those fed on by infected ticks, whereas IgG levels to rS13c5 remained generally low, with only minor differences between groups.

## Discussion

To acquire and transmit tick-borne disease (TBD) pathogens, ticks must feed and circumvent host defense pathways that are primarily orchestrated by serine proteases and tightly regulated by serpins. Tick saliva serpins are therefore regarded as critical regulators of blood feeding and important contributors to the transmission of LD and other TBD pathogens (9, 10, 12). In previous studies, we identified 45 putative serpins in the *Ixodes scapularis* genome (51) and demonstrated that some of these proteins function as tick saliva transmission factors for Bb, the LD pathogen (29). A revision of the *I. scapularis* genome annotation increased the number of encoded serpins to 46 (73). More recently, our reanalysis of the updated genome uncovered 74 serpin-encoding genes located on 8 of the 13 autosomal chromosomes and the unassigned genomic scaffolds. In this analysis, we also refined the classification of tick serpins based on their chromosomal organization (52).

In the present study, we focused on characterizing two distinct serpins, S12c5 and S13c5, which share less than 20% amino acid identity. Both proteins were analyzed together because both are secreted with identical secretion patterns during feeding by uninfected and Bb-infected nymphs (32). Notably, both genes exhibit a similar genomic organization on *I. scapularis* chromosome 5, where they are tandemly arranged, each encoded by a single exon and likely arising from duplication events (52). This conserved genomic organization further supports their designation as infection-responsive tick saliva proteins that contribute to Bb transmission. BLAST analysis revealed that, despite their marked divergence from one another (<20% amino acid identity), S12c5 and S13c5 are highly conserved across other tick species. Amino acid identity with homologous proteins ranges from 60–99%, whereas the functional RCL domain is 80–100% conserved. This level of evolutionary conservation suggests that S12c5, S13c5, and their homologs perform essential functions that support both tick feeding and the transmission of TBD agents (50). The long-term goal of our research is to develop a tick antigen–based vaccine to prevent Lyme disease and other TBDs. Our approach is to induce host immunity that disrupts the tick’s ability to evade both innate and acquired immune defenses at the skin, thereby impairing tick feeding and the transmission of TBD agents. Successful tick feeding requires suppression of host defenses such as blood clotting, inflammation and tissue repair, nociception, and complement activation. To gain functional insights, we first profiled the inhibitory activity of yeast-expressed rS12c5 and rS13c5 against a panel of 18 effector serine proteases as well as components of the complement system. Our results indicate that S13c5 is a significantly more potent inhibitor of proteases involved in blood clotting, inflammation/tissue repair, and nociception compared to rS12c5. Although both serpins effectively inhibit complement activation via the lectin pathway and, to a moderate extent, the alternative pathway, molecular modeling data suggest that S13c5 functions as a more efficient inhibitor of complement activation overall.

Although not empirically quantified, we expect that the tick injects miniscule amounts of native S12c5 and S13c5 into the feeding site to block functions of host defense system proteases. In this scenario, we also expect that the less amount of serpin molecules will block the functions of its target protease under physiological conditions. Thus, functional efficiency of rS13c5 in this study may approximate activities under physiological conditions (SI of 1) (55). With just a few rS13c5 molecules at low SIs,of 2.5, 1.9, and 1.0, respectively blocked proteases involved in blood clotting factor (f) Xa and plasmin and tissue repair (trypsin IV). Remarkably, the rate at which rS13c5 inhibition of plasmin (3.97 x 10^5^M^-1^s^-1^) and trypsin IV (1.08 x 10^5^M^-1^s^-1^) occurs resembles that at which a typical serpin inhibits the activity of its cognate protease (≥10^5^ M^-1^ s^-1^) (55). The slower rate of rS13c5 inhibition of blood clotting factor (f) Xa (8.56 x 10^3^M^-1^s^-1^) may be explained by the need for a yet unknown co-factor such as glycosaminoglycans (GAGs) as observed with other serpins (74–77). The need of a cofactor could also explain the weak inhibitory activity observed with rS12c5.

Both fXa and trypsin IV activate protease-activated receptors (PAR)-1, PAR-2, and PAR-4 (78, 79), thereby contributing to inflammatory and tissue repair responses. Consequently, the inhibitory activity of rS13c5 against these proteases may simultaneously suppress host hemostatic and inflammatory pathways, facilitating tick feeding and promoting *B. burgdorferi* colonization of the vertebrate host. In contrast to fXa, and its importance in the formation of a stable fibrin clot, plasmin is a key component for the degradation of these clots (fibrinolysis) (80) and activation of proinflammatory cells (81). At first glance, the potent inhibition of plasmin by S13c5 appears paradoxical. Because successful tick feeding depends on limiting both blood coagulation and inflammation, inhibition of fibrinolysis would be expected to favor clot persistence and potentially impair bloodmeal acquisition (82). However, the selective secretion of S13c5 by Bb-infected ticks suggests that its anti-plasmin activity may provide a pathogen-specific advantage. One possible explanation involves the role of plasmin in neutrophil recruitment. Plasmin-mediated degradation of fibrin generates fibrin degradation products (FDPs), which function as potent neutrophil chemoattractants (83, 84). Neutrophils are known to entrap and kill Bb through the release of neutrophil extracellular traps (NETs) (85). Thus, although inhibition of plasmin may partially counteract tick-mediated suppression of clotting and inflammation, S13c5 may enhance the survival of transmitted spirochetes at the feeding site by limiting plasmin-dependent neutrophil recruitment. Our findings are also intriguing in the context of Bb pathogenesis. Several borrelial proteins, including BBA70 (86), OspA (87), and OspC (88) bind plasminogen and promote its activation to plasmin (89). Activated plasmin facilitates degradation of extracellular matrix components and contributes to spirochete dissemination within the host (90, 91). One possibility is that S13c5 primarily functions during the early stages of transmission, when evasion of innate immune defenses at the feeding site is more critical than dissemination. Future studies should investigate how S13c5 influences plasmin acquisition by Bb and determine its impact on spirochete dissemination and pathogenesis in vivo.

The complement system represents another key component of host innate immunity that ticks must circumvent to successfully feed and transmit pathogens (92). Previous studies have identified several *I. scapularis* salivary proteins, including Isac (93), Salp14 (94), Salp20 (95), *Ixs*S17 (47), and *Ixs*S41 (96) that inhibit complement activation. Here, we show that both S12c5 and S13c5 are potent inhibitors of complement activation through the MBL pathway and, to a lesser extent, the alternative pathway.

To elucidate the mechanisms underlying S12c5 and S13c5 inhibition of complement, we performed pull-down assays coupled with ELISA, western blotting, LC-MS/MS analysis, and molecular modeling. Structural modeling predicted that S13c5 may interact more favorably than S12c5 with complement-associated proteases. Consistent with this prediction, both proteins bound multiple complement components. In particular, ELISA and western blot analyses demonstrated interactions with MASP-1/3, the effector serine proteases of the lectin pathway, suggesting that inhibition of these proteases contributes to the observed suppression of MBL-mediated complement activation. In addition, both serpins associated with several other protease and non-protease complement components, indicating that their inhibitory activities may involve multiple targets. A notable discrepancy emerged between the immunological and proteomic analyses. Although MASP-1/3 was readily detected in S12c5 and S13c5 pull-down eluates by ELISA and western blotting, neither protein was identified by LC-MS/MS. While low-abundance proteins can escape detection in proteomic analyses because of ion suppression by more abundant proteins (97, 98), this explanation appears unlikely given the relatively high circulating concentrations of both MASP-1 and MASP-3 (99, 100). Furthermore, because MASP-1 and MASP-3 contain multiple trypsin cleavage sites, inefficient proteolytic digestion is also an unlikely explanation. The basis for this discrepancy therefore remains unclear and warrants further investigation.

Although neither rS12c5 nor rS13c5 inhibited activation of the classical pathway, both pull-down eluates contained substantial amounts of C1q, C1r, and C1s, the components of the C1 complex that initiates classical pathway activation (101, 102). The biological significance of these interactions remains uncertain, particularly given the lack of measurable inhibition of classical pathway activity. In contrast, both serpins also associated with properdin and factor H, key regulators of the alternative pathway. Properdin stabilizes the alternative pathway C3 convertase and promotes complement activation (103, 104), whereas factor H negatively regulates the pathway by serving as a cofactor for factor I-mediated cleavage of C3b (105). These interactions may contribute to the moderate inhibition of the alternative pathway observed for both proteins and further support the notion that S12c5 and S13c5 target multiple components of the complement cascade.

Serpins are classically recognized for their ability to inhibit serine and cysteine proteases (40). Consistent with this function, our pull-down analyses identified multiple complement proteases together with their endogenous regulatory serpin, C1-inhibitor. In addition, several non-protease complement components, including C4b-binding protein, C3, C4, C4a, C5, C6, C8, and C9, were detected in S12c5 and S13c5 eluates. Similar patterns were observed for other proteolytic systems. For example, thrombin, plasma kallikrein, and plasmin were recovered together with their respective regulatory serpins, antithrombin, kallistatin, and α2-antiplasmin. One possible explanation is that initial interactions between S12c5 or S13c5 and target proteases facilitated the co-purification of additional proteins associated with the same regulatory pathways. Similar observations were reported in our previous characterization of IxsS17 (renamed S61c10) (47). However, whether these interactions reflect physiologically relevant complexes formed in vivo remains to be determined.

Consistent with our protease inhibitor profiling data, S12c5 showed no effect on plasma clotting. In contrast, S13c5, an effective inhibitor of blood clotting factor Xa (fXa), prolonged plasma clotting time in the recalcification assay but not in the activated partial thromboplastin time (aPTT) assays, which assess the integrity of the intrinsic coagulation pathways (106, 107). These findings suggest that S13c5 exerts its anticoagulant activity through inhibition of fXa within the common coagulation pathway. Because fXa is the central protease responsible for thrombin generation and subsequent fibrin clot formation, its inhibition would be expected to impair clot development and stabilization (108). Interestingly, although S13c5 inhibited fXa activity in substrate hydrolysis assays, fXa was not detected in the corresponding pull-down eluates. Conversely, despite lacking measurable anticoagulant activity, S12c5 co-purified with several coagulation-related proteins, including thrombin, factor XIa, factor XIIa, and factor V. Together, these observations underscore the complexity of interpreting pull-down interactions and suggest that protein association alone may not reliably predict functional inhibition.

The MBL pathway plays an important role in host defense against Bb (109, 110). Consistent with their strong anti-complement activity, both rS12c5 and rS13c5 significantly protected Bb from serum complement–mediated killing. Moreover, co-inoculation of S13c5 enhanced Bb colonization of murine tissues, supporting a role for this serpin in promoting pathogen survival during the early stages of infection. Because our long-term objective is to identify tick antigens suitable for vaccine development against Lyme disease and other tick-borne diseases, we also evaluated the immunogenicity of both proteins using sera from rabbits exposed to uninfected or Bb-infected *I. scapularis* nymphs (24, 96). Notably, S13c5, which displayed the broader and more potent inhibitory activity against innate immune proteases, elicited weaker antibody recognition than S12c5. Although the basis for this difference remains unknown, reduced immunogenicity could represent an adaptation that preserves the function of critical salivary proteins involved in immune evasion.

In summary, our findings identify S12c5 and S13c5 as infection-responsive tick salivary serpins that contribute to the modulation of host innate immune defenses during tick feeding and pathogen transmission. While S13c5 exhibited potent inhibitory activity against multiple innate immune proteases and enhanced Bb survival, S12c5 showed limited activity against the proteases tested despite sharing a similar expression profile and genomic organization. One possibility is that S12c5 serves a complementary or supportive role that remains to be defined. Alternatively, it may function as an immunological decoy, diverting host antibody responses away from more functionally important salivary proteins such as S13c5. Further studies will be required to distinguish between these possibilities. Collectively, these findings reinforce the concept that tick salivary proteins act cooperatively to suppress host immune defenses and facilitate pathogen transmission. In addition, the potent inhibition of fXa by S13c5 and its measurable anticoagulant activity suggest that this protein may warrant further investigation as a source of novel anticoagulant molecules with potential therapeutic applications for the treatment of thromboembolic disorders such as deep vein thrombosis and pulmonary embolism, and possibly atherosclerosis (108, 111, 112). Our results highlight the dual significance of these proteins as potential targets for tick antigen-based vaccines to prevent LD and as promising candidates for further exploration of their therapeutic applications.

## Materials and Methods

### Ethics Statement

The described experimental procedures were executed following the animal use protocol approved by Texas A&M University Institutional Animal Care and Use Committee (IACUC) (AUP 2018-001 and 2020-0089) to meet all federal requirements, as defined in the Animal Welfare Act (AWA), the Public Health Service Policy (PHS), and the Humane Care and Use of Laboratory Animals.

### Cloning of I. scapularis serpin (S) 12c5 and S13c5

S12c5 and S13c5 were originally identified among proteins secreted at high abundance by Bb-infected *I. scapularis* nymphs (24). Template cDNA from 72 h rabbit fed Bb*-*infected nymphs was used for 3’ prime rapid amplification of cDNA ends (RACE) using the SMARTer Clontech 3 and 5’ race kit (Takara Bio, San Jose, CA, USA) as previously described (96). Primers that were used with the company provided universal amplification primer (UAP) were designed using the amino acid sequences to S12c5: XM_040215702.3 and S13c5: XM_029978268.4 obtained from GenBank (S13c5-Forward: ^5’^ATGTGGTTCCCGGCGCTCCTG^3’^ and S12c5-Forward: ^5’^ATGAAACGCTGCACCCTGGTC^3’^). Subsequently, a nested PCR was done using forward primers and S13c5-Reverse: ^5’^TCACAAGTCCAGCACTCGG^3’^ and S12c5-Reverse: ^5’^TCACACCTCTTGAAGCCTTCCG^3’^. PCR amplicons were cloned and sequenced using standard methods. The confirmed sequences were used in comparative sequence analysis and recombinant protein expression (below).

### Comparative sequence analysis

For comparative sequence analysis, pairwise alignment in MacVector was used to compare amino acid sequences of S12c5 (XP_040071636.1) and S13c5 (XP_029834128.2). Multiple sequence alignment tool in MacVector was used to gauge the relationship of S12c5 and S13c5 protein sequences to their orthologs downloaded from GenBank. S12c5 was compared to *Ixodes persulcatus* (KAG0414789.1), *I. hexagonus* (CAN7997772.1), *Amblyomma americanum* (KAK8787056.1), *Dermacentor silvarum* (XP_049517673.1), *D. andersoni* (XP_050043950.1), *Rhipicephalus sanguineus* (XP_037510184.1), *R. appendiculatus* (KAL1425160.1 and KAL1471163.1), *Hyalomma asiaticum* (KAH6937721.1), *Haemaphysalis longicornis* (KAH9371121.1), *R. microplus* (KAH8009656.1), and the soft tick, *Ornithodorous turicata* (XP_064479489.1). Similarly, S13c5 (XP_029834128.2) was compared to *I. persulcatus* (KAG0414788.1), *Amblyomma americanum* (JAI08960), *D. andersoni* (XP_050043949.1), *Dermacentor silvarum* (XP_049517673.1), *H. longicornis* (KAH9371123.1), *R. haemaphysaloides* (QNV48485.1), *R. microplus* (XP_037285924.1), and *R. sanguineus* (XP_037510183.1).

### Expression and purification of wild-type and RCL-deletion mutant recombinant (r) S12c5 and S13c5

Synthesis of expression constructs for wildtype recombinant mature protein in pPICZαA plasmid with a hexa-histidine tag (His-tag) was outsourced (Biomatik USA, Cambridge, Canada). Expression constructs for reactive center loop deletion mutants (or truncated) were synthesized in the lab using Biomatik constructs as template. Subsequently, *Pichia pastoris* X-33 cells were transformed by electroporation as previously described (23, 48, 96). Pilot expression (10 mL cultures) of recombinant proteins was induced for five days by adding 1% methanol (final concentration). Expression of both proteins was confirmed by western blot using mouse THE^TM^ His Tag Antibody (HRP [GenScript Biotech, Piscataway, NJ, USA]) 1:5000-diluted (0.002 µg). Subsequently, large scale (1 L) recombinants were done as optimized during pilot expression. The pPICZαA plasmid is optimized to secrete recombinant proteins into spent media. Routinely, recombinant proteins were precipitated out using ammonium sulfate saturation. Precipitants were resuspended, dialyzed against affinity column binding buffer (20 mM Tris-HCl, 150 mM NaCl, 5mM imidazole, pH 7.4) and filtered with a 0.2 µm filter (VWR, Radnor, PA, USA). Purification was done using HiTrap Chelating HP columns (Cytiva Life Sciences, Marlborough, MA, USA) and Tris-HCl, imidazole (23, 48). Routinely, standard SDS-PAGE and silver staining along with western blotting analysis using the antibody to the His-fusion tag were used to verify expression and affinity purification of recombinant proteins. Spin columns with a 30 kDa cut-off (Pall Corporation, Port Washington, NY, USA) were used for volume reduction and buffer exchange the substrate hydrolysis reaction buffer (20mM Tris-HCl, 150mM NaCl, pH 7.4). Protein concentration was verified using the Pierce BCA Protein Assay Kit (ThermoFisher Scientific, Waltham, MA, USA).

### Validation of glycosylation

Affinity purified rS12c5 and rS13c5 protein bands appeared diffuse suggesting glycosylation. To verify, approximately 100 µg of each protein was subjected to deglycosylation for the removal of both N- and O- linked glycans under denaturing conditions for 1h at 37° C. The NEB Protein Deglycosylation Mix II (NEB, Inswic, MA, USA) was used, following manufacturer instructions. Approximately 1µg of untreated and treated rS12c5 and rS13c5 were resolved on a 7.5% native gel and subjected to silver staining using the Pierce Silver Stain kit (ThermoFisher Scientific) to verify deglycosylation.

### Protease inhibitory activity profiling

Given the role of tick secreted serine protease inhibitors and their importance in blocking host proteases involved in protection against tick feeding, the inhibitory activity of rS12c5 and rS13c5 was verified by substrate hydrolysis as optimized by our group (23, 48, 55, 96). Routinely, 1 µM of affinity purified rS12c5 or rS13c5 in reaction buffer (20 mM Tris-HCl, 50 mM NaCl, Tween^TM^-20 0.1%, pH7.4) was pre-incubated with different mammalian serine proteases for 15 min at 37 °C. A chromogenic substrate specific to each protease was added at 200 µM final concentration (100 µl total reaction) and its hydrolysis was recorded (A_405nm_) every 11 s for 15 min at 30 °C using the Synergy H1 plate reader (BioTek Instruments Inc, Winooski, VT, USA).

### Stoichiometry of inhibition (SI)

The SI of rS13c5 against fXa, fXla, plasmin, pancreatic trypsin, and trypsin IV, proteases whose activity was inhibited by ≥80% was determined as optimized by our group (23, 33, 96). Different amounts of rS13c5 were incubated with constant amounts of each protease (11.84 nM of faXa, 6.66 nM of fXla, 9.88 nM of plasmin, 1.2 nM of pancreatic trypsin, and 3.74 nM of trypsin IV) for 1 h at 37 °C. Thereafter, colorimetric substrates were added, and the residual enzymatic activity was determined using the Synergy H1 plate reader (A_405nm_). These data are presented as residual activity (V_i_/V_0_) against rS13c5 to protease molar ratio as previously described (23, 33, 96).

### Rate of inhibitory reactions (k_a_)

Following a preliminary assessment of rS13c5 inhibition of plasmin, trypsin IV, and factor Xa as previously described (55), we concluded that the protein is a fast inhibitor of plasmin and trypsin IV. Based on this information, the continuous method for plasmin and trypsin IV, and the discontinuous method for fXa were conducted to determine the rate of inhibitory reaction (k’). Under the discontinuous method, increasing amounts of rS13c5 (2.9-95.0 nm) were pre-incubated with 9.5 nM of factor Xa for 0-15 min at 37 °C and the OD_450nm_ recorded. The k_obs_ was obtained as published (23, 33, 96) and best fit line of k_obs_ values were plotted against different amounts of rS13c5 that were used to obtain the second-order rate constant k_a_ (23, 48, 55).

Under the continuous method, increasing amounts of rS13c5 were mixed with constant amounts of protease (16.0-60.0 nM of r*Ixs*S19 with 1.6 nM of plasmin; 0.9-22.4 nM of rS13c5 with 3.74 nM of Trypsin IV). The serpin-protease mix was added to a 96-well plate containing their corresponding chromogenic substrates (0.2 mM) and OD_405nm_ was recorded with a Synergy H1 microplate reader every 19 s for 30 min. The observed pseudo-first order rate constant k_obs_ were derived from exponential fit of progress curves using nonlinear regression analysis to calculate the apparent first order rate constant k_obs_. The apparent second order association rate constant, k’, was obtained from slopes of linear regression analysis of rS13c5 concentration against k_obs_ values, accounting for K_M_ of plasmin (3 x 10^-4^mol/L) and trypsin IV (4.1 x 10^-6^ mol/L), and SI as previously described (55).

### Effect of rS12c5 and rS13c5 on blood clotting

Blood clotting assays were done using the recalcification time (holistic assessment of blood clotting) as described (33, 113). Various amounts of rS12c5 or rS13c5 (0.28, 0.50, 1.12, 2.50, and 4.50 µM) in 40 µL of Tris-HCl buffer were gently mixed with universal coagulation reference plasma (UCRP) (ThermoFisher Scientific) and incubated for 15 min at 37 °C. Immediately after incubation, 10 µL of pre-warmed (37 °C) CaCl_2_ (150 mM) was used to initiate plasma clotting which was monitored (A_650nm_) every 20s for 16 min with a Synergy H1 microplate reader. To estimate plasma clotting time (X axis value), optical density (*A*_450 nm_) data were fitted to a simple linear regression model in PRISM 10, with the Y axis value (clotting OD) set at 0.01. The impact on clot quality was assessed as previously described, with modifications (58).To evaluate the firmness or overall quality of the plasma clot, A450 nm data were further analyzed using a one-phase decay model in PRISM 10 to determine the plateau. These values were then used to calculate the percentage (%) effect of rS12c5 and rS13c5 on clot.

### Effects of rS12c5 and rS13c5 against complement activation

The WiesLab Complement System kit (Svar Life Science AB, Malmö, Sweden) was used to independently evaluate the effects of rS12c5 and rS13c5 on each of the three complement activation pathways: classical, alternative, and MBL as optimized by our lab (47, 96, 114). Approximately 4 µM of rS12c5 or rS13c5 were gently mixed and pre-incubated with the company provided human serum (positive control, PC) for 30 min at 37 °C. The reaction was repeated to evaluate RCL deletion mutants. The positive control was also incubated with buffer control (20 mM Tris-HCl, 150 mM NaCl, pH 7.4). Subsequently, the pre-incubated components and the company-provided negative control (inactive serum) were transferred onto the company-provided wells for an additional incubation at 37 °C for 60 min. The conjugate and substrate were added for 30 min each, following manufacturer instructions. A Synergy H1 plate reader was used to obtain *A_405nm,_* and deposition of the membrane attack complex (MAC) was calculated using the formula suggested by the manufacturer: 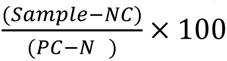

Preliminary findings showed that excess amounts (4 µM) of rS12c5 and rS13c5 significantly reduced the deposition of MAC via the MBL pathway. Thus, the assay was repeated using five-fold serially diluted rS12c5 and rS13c5 (4-0.0064 µM).

### Pull-down assay to elucidate the mechanisms by which rS12c5 and rS13c5 inhibit complement activation

To investigate the mechanisms by which rS12c5 and rS13c5 inhibited complement activation, pull-down magnetic beads (ThermoFisher Scientific) were used to perform a pull-down assay as published (114). Briefly, purified His-tagged rS12c5 or rS13c5 (10 µg) in binding buffer (700 µL; 50 mM sodium phosphate, pH 8.0, 300 mM NaCl, 0.01% Tween^TM^-20) was coated onto magnetic Dynabeads™ (2 mg). Subsequently, beads were washed with washing buffer (100 mM sodium phosphate, 600 mM NaCl, 0.02% Tween-20, pH 8.0) to remove non-specifically bound proteins. HCS (human complement serum; Innovative Research, Inc., Novi, MI, USA) diluted to 10% in the pull-down buffer (6.5 mM sodium phosphate 140 mM NaCl, 0.02% Tween-20; pH 7.4) was incubated with the bead–bait complex for 2 h with mixing at room temperature (RT). Non-bound proteins were discarded, and beads were washed 4 times with the wash buffer (100 mM sodium phosphate 600 mM NaCl, 0.02% Tween-20; pH 8.0). Following appropriate washing, bound protein complexes were eluted by incubating the beads with elution buffer (300 mM Imidazole, 50 mM sodium phosphate 300 mM NaCl, 0.01% Tween-20: pH 8.0) for 5 min at RT. For control beads and HCS only incubation was processed and eluted as above.

Subsequently, rS12c5 and rS13c5 pull down eluates along with negative control eluate (HCS and beads only incubation) and positive control (1% HCS) were subjected to standard western blotting analysis using antibodies to complement system proteins. Primary antibodies included the HRP conjugated His Tag Antibody (GenScript) mouse anti-MASP1/3, anti-activated C3 (Santa Cruz Biotechnology, Inc., TX, USA), goat anti-human C3, C5, and C9 (Complement Technology, Inc., Tyler, TX, USA). HRP conjugated anti-mouse (SouthernBiotech) and rabbit anti-goat IgG-HRP (Invitrogen, Waltham, MA, USA) were used as secondary antibodies and chemiluminescence was detected and visualized as described before(49).

For ELISA analysis, 96-well plates (ThermoFisher Scientific) were coated (in duplicate) with 100 µL containing 200 ng of rS12c5, rS12c5/HCS eluate, rS13c5, rS13c5/HCS eluate, positive control (1% HCS) and bead and HCS only eluate (40 ng, empirically determined for minimal non-specific binding). Following blocking and washing as described before (24, 47, 49), primary and secondary antibodies were used at the same concentration used for western blots. The 1-Step Ultra TMB-ELISA substrate and 2N sulfuric acid were added, and A_405nm_ was measured with a Synergy H1 plate reader.

### LC–MS/MS Identification of Human Complement Serum Proteins Interacting with rS12c5 and rS13c5

LC-MS/MS analysis was done as published (114). Briefly, in-solution protein digestion using trypsin was used to prepare pull down eluates (10 µg) for LC-MS/MS analysis. Eluate proteins were digested overnight at 37°C with trypsin at a 50:1 ratio (protein-to-trypsin) and submitted for LC-MS/MS analysis. LC-MS/MS analysis was performed on the Orbitrap Fusion Tribrid Mass Spectrometer (ThermoFisher Scientific) equipped with a Dionex UltiMate 3000 reverse-phase nano-UHPLC system (ThermoFisher Scientific). Mass spectrometry data were acquired in positive mode at a resolution of 120,000 (at *m*/*z* 200) in the *m*/*z* range of 400–1600. Proteome Discoverer 2.4 software (ThermoFisher Scientific) was used to analyze the MS/MS spectra of peptide ions to search against the database of human plasma proteins downloaded from GenBank (www.ncbi.nlm.nih.gov) combined with S12c5 and S13c5 protein sequences was searched using tandem mass spectra to identify proteins as published (114). Subsequently, the pathways that were enriched in interactions between rS12c5 or rS13c5 and plasma proteins were identified using the Reactome database (63). Accession numbers of plasma protein interactors with rS12c5 or rS13c5 were loaded onto the Reactome database server and highly significant pathways were reported.

### Molecular modeling

The *Ixodes scapularis* leukocyte elastase inhibitor A (S13c5) (NCBI Reference Sequence: XP_029834128.2), corresponding to residues 18–390, and the *I. scapularis* intracellular coagulation inhibitor 1 (rS12c5) (NCBI Reference Sequence: XP_040071636.1), corresponding to residues 19–399, were modeled in complex with residues (res) corresponding to the Peptidase S1 (SP) domains of C1s (UNIPROT#: P09871, res 438-680), C1r (UNIPROT#: P00736, res 464-702), MASP-1(UNIPROT#: P48740, res 449-696), MASP-2 (UNIPROT#: O00187, res 445-684), C2 (UNIPROT#: P06681, res 464-744), FB (P00751, res 477-757), FD (P00746, res 26-253), and FI (UNIPROT#: P05156, 340-574) using AlphaFold Server (115). Structural alignment between the AlphaFold3-predicted C1s/S19 or C1s/S20 model and the crystal structure of the C1s/C1-INH (PDB, 8W18, (68) complex was performed using the *super* function in PyMOL v 2.4.1 (The PyMOL Molecular Graphics System, Schrödinger, LLC).

### Serum complement rescue assay

The importance of the complement cascade in clearance of Bb is established (109, 116–118). Thus, we sought to evaluate the effects of rS12c5 and rS13c5, on the complement sensitive Bb strain B314 (69, 118). and resistant (“BmA”) Bb strains via a serum complement sensitivity assay. For this assay, complement sensitive, “B314”, represents B314/pBBE22*luc* as previously described (49, 119). On the other hand, complement resistant, “BmA”, represents B314 with the BBK32-like protein to *Borrelia miyamotoi* FbpA, cloned into pBBE22*luc* vector (69). BmA has been identified as a human C1r inhibitor (69, 120). Strain B314 represents B314/pBBE22*luc* as previously described (49, 119). “BmA”, represents a BBK32-like protein from *Borrelia miyamotoi* FbpA that exhibits high-affinity binding and inhibition to human C1r and serves a positive control for the serum rescue assay (69, 120). The “BmA DA” derivative is a double alanine mutant that does not bind or inhibit human C1r and thus serves as a negative control. Both BmA samples were incubated in normal human serum at 0.25 μM as describe below and were compared relative to untreated normal human serum, which effectively kills the serum sensitive strain B314 derivative. In a sterile microtiter plate, either 4 µM (≤10 µl as total volume) of rS12c5 or rS13c5, as well as heat inactivated versions of both S12c5 and S13c5 and RCL truncated forms, were incubated with 15 µL of normal human serum (Complement Technology) in a 96-well plate for 30 min at 37 °C. Subsequently, 85 µL of complement sensitive B314 was diluted to 1x10^6^ spirochetes/ml and added to its respective well to a final concentration of 15% normal human serum and incubated for 90 min at 32 °C under constant shaking at 100 rpm. After incubation, a dark field microscope was used to score spirochete survival on ten fields based on mobility and unimpaired membrane as before (96, 116, 118).

### Effect or rS12c5 on Bb colonization of mice

Given the inhibitory activity of rS13c5 against complement activation and proteases involved host innate immune responses against tick feeding and Bb transmission and dissemination, the effects of co-inoculating rS13c5 and Bb on spirochete colonization of mouse organs was evaluated. Twelve female C3H/HeN mice (9-10 weeks) were purchased from Envigo (Indianapolis, IN, USA) and allowed to acclimate for seven days. Following this period, they were anesthetized with 3% isoflurane through a nose cone (600 cc of O_2_) to shave and cleanse the scapular area as previously described (96). The prepared area was intradermally injected with: (A) 1 x 10^6^ spirochetes, (B) 1 x 10^4^ spirochetes, (C) 1 x 10^6^ spirochetes and 25 µg of rS13c5, or (D) 1 x 10^4^ spirochetes and 25 µg of rS13c5 (three mice per treatment). Additionally, the tibiotarsal joints of both feet were measured prior to needle inoculation, as well as seven and fourteen days after inoculation. Mice were euthanized fourteen days after inoculation in a CO_2_ chamber and cervical dislocation, immediately followed by organ collection (ear skin, tibiotarsal joint, heart, dura mater, bladder, spleen, liver, and inguinal lymph node) for gDNA extraction using the DNeasy Blood & Tissue Kit (Qiagen, Germantown, MD, USA). These gDNA extractions were used for quantitative (q) PCR with murine β- actin (F: 5’ CAAGTCATCACTATTGGCAACGA 3’ and R: 5’ CCAAGAAGGAAGGCTGGAAAA 3’) and Bb *flaB* (F: 5’ TCTTTTCTCTGGTGAGGGAGCT 3’ and 5’ TCCTTCCTGTTGAACACCCTCT 3’) primers as previous described (121) (96) (with iTaq Universal SYBR Green Supermix (BioRad Laboratories, Ann Arbor, MI, USA) in 50 µL reactions. Additionally, mouse (316.8 ng/µl) and Bb (0.300 ng/µl) gDNA (5-fold diluted) were used as known standards. The cycling conditions were: 50° C for 2 minutes (min), activation at 95° C for 10 min, and 40 cycles of denaturation at 95° C for 15 seconds (s) and annealing/extension at 60° C for 1 min. Data was analyzed to calculate the relative Bb *flaB* gene expression in the organs collected using the delta delta Ct method (2^-ΔΔCt^) as published (47, 49). The spirochetes used for needle inoculation of C3H/HeN mice were Bb strain 31 MSK5 cultured until mid-log phase in BSK-II media at 32° C with 1% CO_2_.

### Immunogenicity analysis using standard western blotting and ELISA analysis

To assess the immunogenicity, various amounts (0.5 and 1 µg) of glycosylated (DG^-^) and deglycosylated (DG^+^) S12c5 and S13c5 were resolved on 12% SDS-PAGE gels and subjected to standard western blotting as described (96). PVDF membranes (Sigma-Aldrich, Burlington, MA, USA) were blocked (skim milk [5%] in phosphate buffered saline-Tween^TM^-20, 0.05% [PBST]) for 1 h at RT. Thereafter, membranes were incubated with empirically determined 16 µg of NZW rabbit purified IgG or 1:100-diluted serum (for IgM) collected before and after being infested with uninfected or Bb*-*infected nymphs overnight at 4 °C. The IgM sera used in this study was collected one week after tick feeding and purified IgG was obtained from sera collected two weeks after tick feeding. Subsequently, membranes were washed with PBST and incubated with 1:5000-diluted goat anti-rabbit HRP conjugated IgG (SouthernBiotech, Birmingham, AL, USA) for 1h at RT. Thereafter, membranes were washed as before, and antibody binding was detected by chemiluminescence using the SuperSignal West Dura Extended Duration Substrate (ThermoFisher Scientific) and visualized using the ChemiDoc MP imaging system (BioRad Laboratories)

The immunogenicity of both serpins was further confirmed by ELISA. Routinely, glycosylated (DG^-^) and deglycosylated (DG^+^) S12c5 and S13c5 (1 µg) were five-fold serially diluted (1-0.25 g) and used to coat a 96-well plate (ThermoFisher Scientific) overnight at 4° C. Following washing with PBST and blocking of wells (with blocking buffer for 1 h at room temperature), purified IgG 16 µg was two-fold serially diluted (16-2 µg) and incubated with coated rS12c5 and rS13c5 overnight at 4 °C. Subsequently, plates were washed and incubated with 1:2000-diluted goat anti-rabbit IgG or HRP conjugated IgM (SouthernBiotech) for 1h at RT. Following washing, 1-Step Ultra TMB-ELISA substrate added to develop the signal and 2N sulfuric acid to halt the reaction. The color intensity (signal) was quantified with a Synergy H1 plate reader (BioTek Instruments Inc.) at A_405nm_.

### Statistical Analysis

Statistical analyses were conducted using GraphPad Prism 10 (GraphPad Software Inc., San Diego, CA, USA). Data are presented as mean ± SEM, with significance set at *p* ≤ 0.05. Analyses included Student’s *t*-test, one-way ANOVA, and ordinary two-way ANOVA.

## Credit authorship contribution statement

**Emily Bencosme-Cuevas:** Conceptualization, Data curation, Formal analysis, Investigation, Methodology, Validation, Visualization, Writing – original draft, Writing – review and editing. **Tae K. Kim:** Investigation, Methodology, Validation, Writing – review and editing. **Thu-Thuy Nguyen:** Data curation, Formal analysis, Investigation, Methodology, Validation, Writing – review and editing. **Moiz A. Ansari:** Data curation, Formal analysis, Investigation, Methodology, Validation, Writing – review and editing. **Shahbaz M. Khan:** Data curation, Formal analysis, Investigation, Methodology, Validation, Writing – review and editing. **Payton G. Smith:** Methodology. **Kyla R. Herr:** Data curation, Formal analysis, Investigation, Methodology, Validation. **Yungui Guo:** Data curation, Formal analysis, Investigation, Methodology, Validation. **William H. Witola**: Data curation, Formal analysis, Methodology, Validation, Resources, Software, Writing – review and editing. **Jon T. Skare:** Data curation, Formal analysis, Funding acquisition, Methodology, Resources, Validation, Writing – review and editing. **Brandon L. Garcia:** Data curation, Formal analysis, Funding acquisition, Methodology, Resources, Software, Validation, Writing – review and editing. **Klaudia I. Kocurek:** Data curation, Formal analysis, Investigation, Methodology, Resources, Software, Validation. **L. Garry Adams:** Conceptualization, Validation, Writing – review and editing. **Stefan H.E. Kaufmann:** Conceptualization, Validation, Writing – review and editing. **Albert Mulenga:** Conceptualization, Data curation, Formal analysis, Funding acquisition, Methodology, Project administration, Resources, Software, Supervision, Validation, Visualization, Writing – original draft, Writing – review and editing.

## Acknowledgments

Studies were supported by NIH grants to AM (AI093858, AI074789, AI138129, and AI119873), and to both JTS and BLG (AI133367 and AI146930). Additionally, we would like to thank the Hagler Institute for Advanced Study at Texas A&M University for supporting EBC through the 2020-2021 HEEP Graduate Fellowship under guidance by SHEK, class of 2019 Hagler fellow. We are grateful to the Chemistry Mass Spectrometry Facility and KIK (TAMU Department of Chemistry) for helping with LC-MS/MS analysis and submission. While most of the work was done at Texas A&M University, EBC is now funded by the Intramural Research Training Award at the NIH. Graphical abstract was created in BioRender. Bencosme Cuevas, E. (2026) https://BioRender.com/3dudhmj.

## Data availability

The datasets and computer code produced in this study are available in the following database: Protein interaction LC-MS/S: MassIVE repository: Dataset Identifier PXD074437 (https://massive.ucsd.edu/ProteoSAFe/private-dataset.jsp?task=6cabffaab25446a0a6b4a057a124335a)

---

**Supplementary Figure S1.**
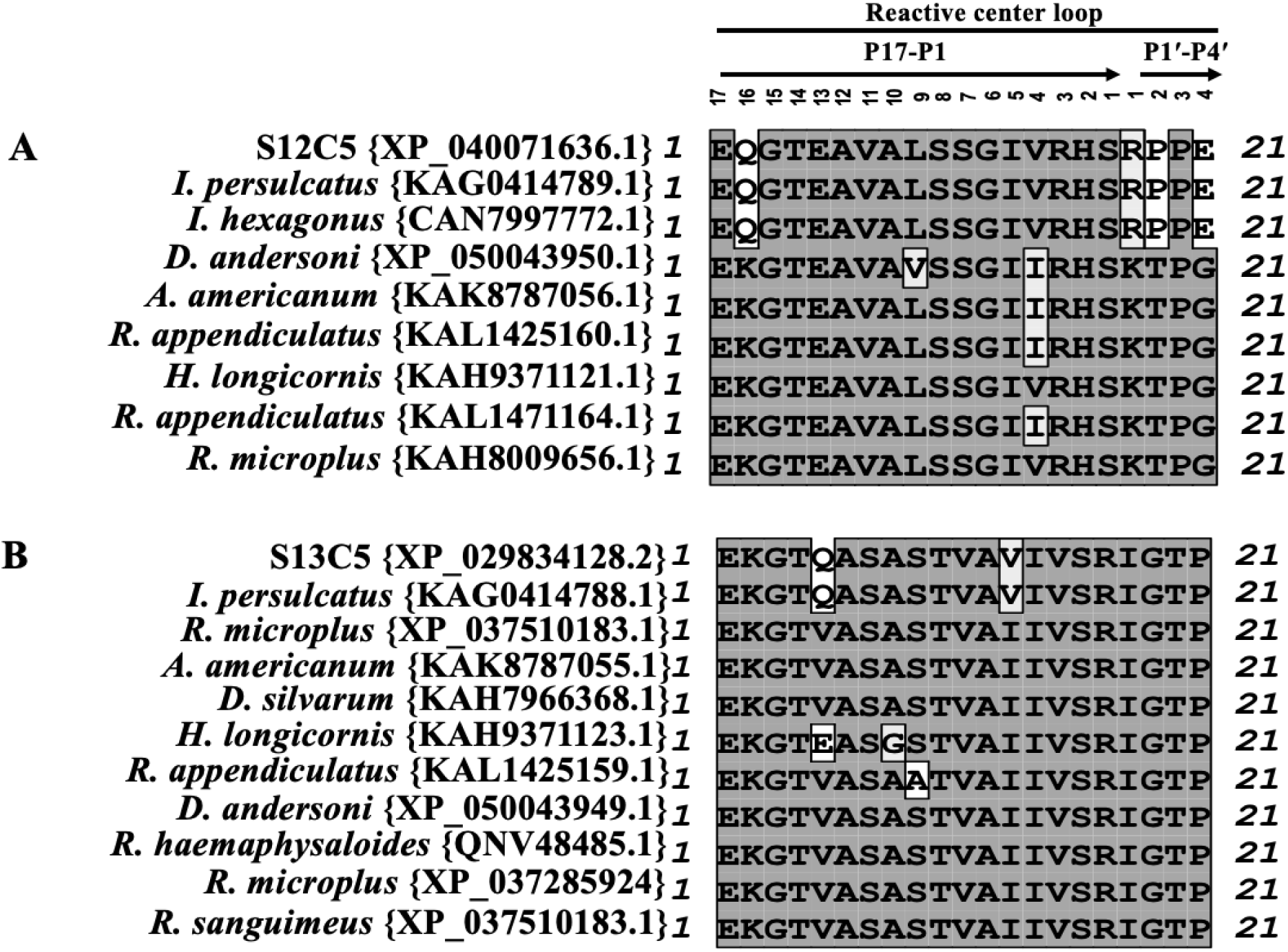
Reactive Center Loop (RCL) amino acid sequences of S12c5 and S13c5. Amino acid sequences from other tick serpins with ≥75% overall amino acid identity to S12c5 and S13c5 were downloaded from GenBank. The RCL region of all amino acid sequences were identified by searching the Conserved Domain Database (CDD) by the National Center for Biotechnology Information (NCBI). RCL sequences were aligned in MacVector to highlight the differences in (A) S12c5 and (B) S13c5 compared to other tick serpins.

**Supplementary Figure S2.**
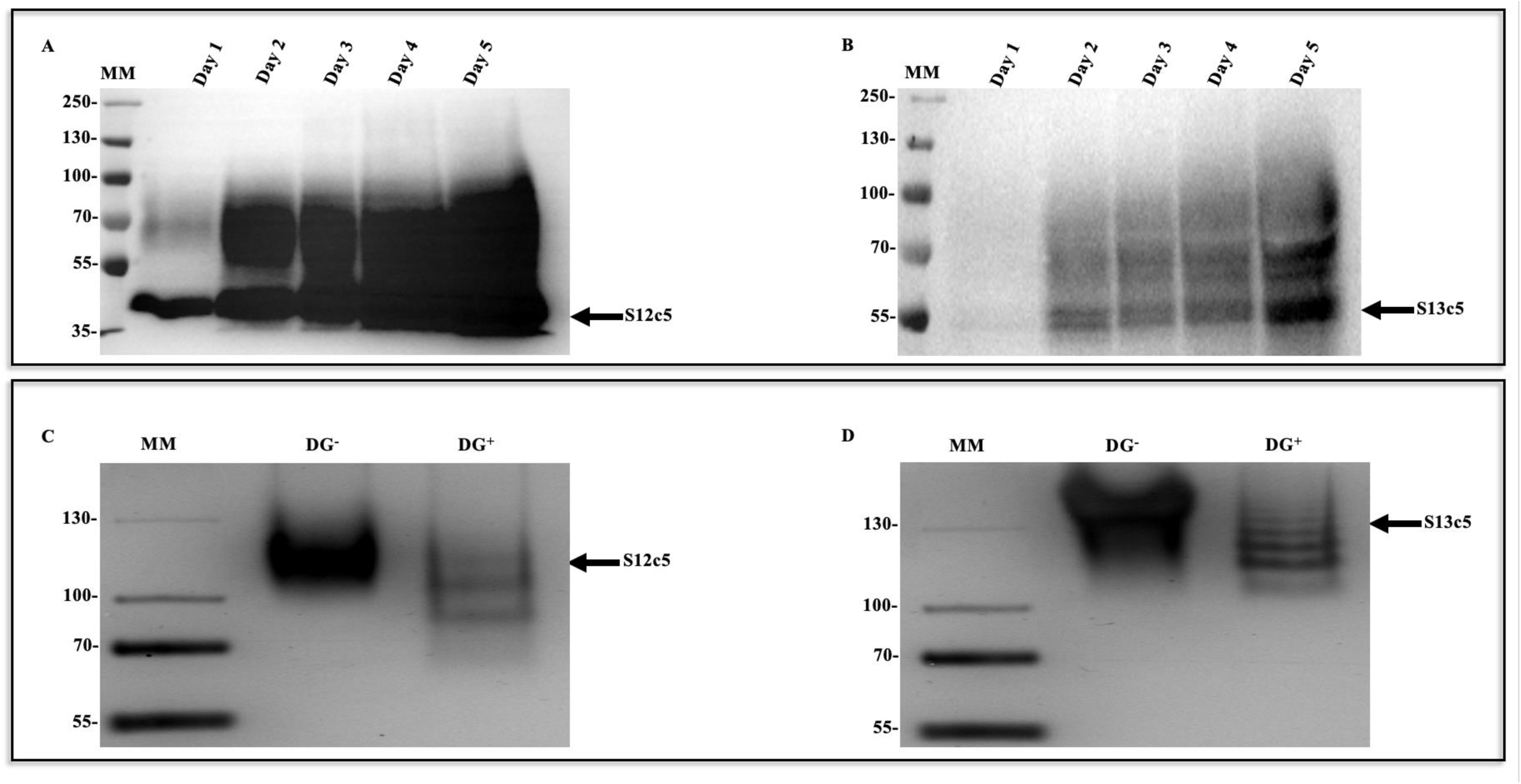
S12c5 and S13c5 are recombinantly expressed as glycoproteins in *Pichia pastoris*. Custom synthesized recombinant (r) protein expression constructs were transformed into *Pichia pastoris* X-33 strain yeast cells by standard electroporation methods and subsequently, adding methanol to cultures at 1% final concentration induced expression of C-terminus histidine tagged rS12c5 (A) and rS13c5 (B) for five days. Expression of each protein was verified by western blot using 1:5000-diluted antibodies to the histidine tag, and affinity purified. Given that both proteins appear are glycosylated as suggested by diffuse banding pattern, approximately 100 µg of each protein was incubated with the deglycosylation enzyme mix for 1 h at 37 °C. To confirm complete deglycosylation, approximately 1 µg of untreated (DG^-^) and deglycosylation enzyme mix treated protein (DG^+^) were resolved on a 7.5% native gel and subjected to silver staining. The untreated and treated *r*S12c5 (C) and rS13c5 (D) are marked by arrowheads.

**Supplementary Figure S3.**
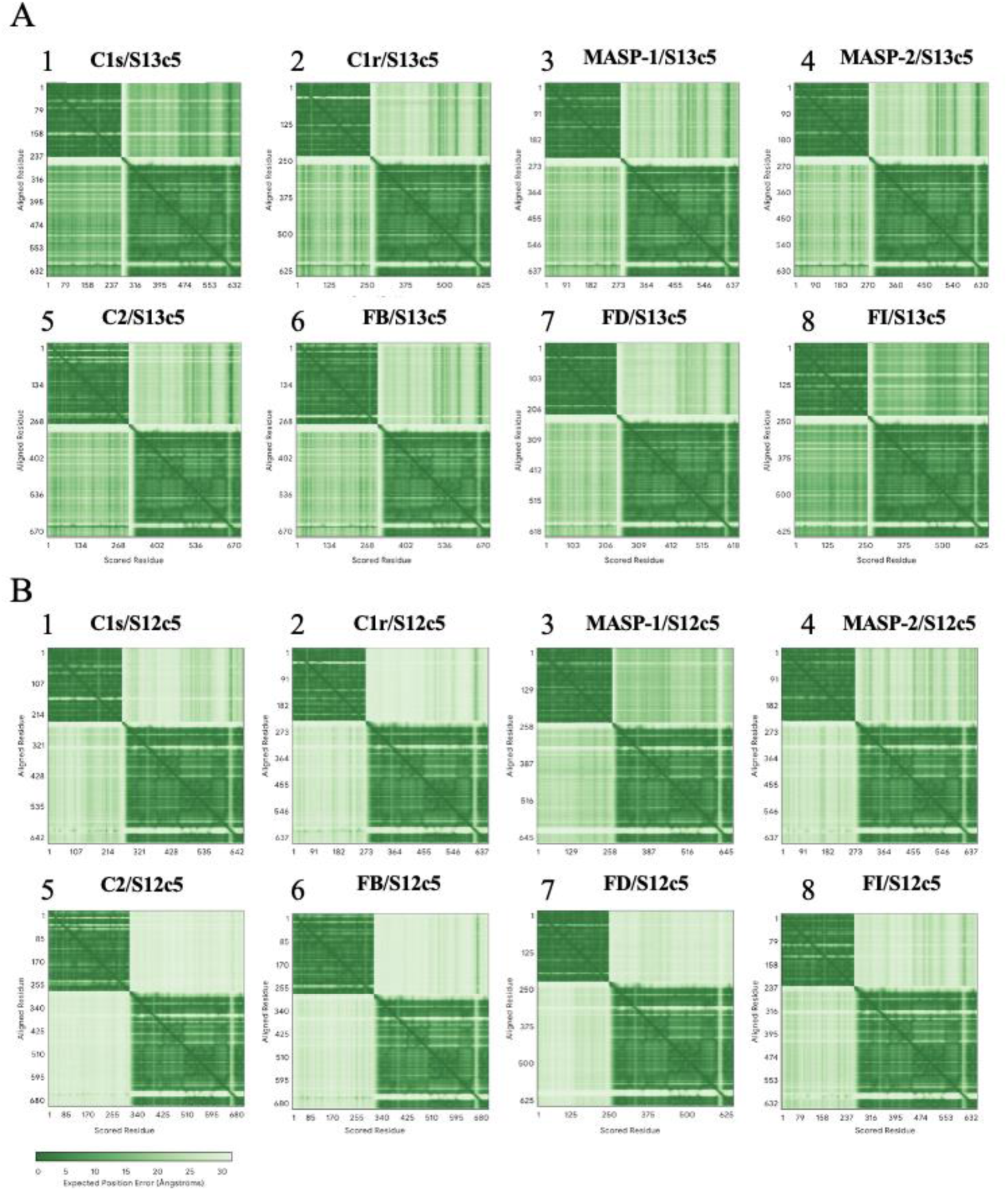
Assessment of AlphaFold3 model quality. (A) The predicted aligned error (PAE) plot is shown for each S13c5 model. In each plot S13c5 corresponds to residues 18-390. For individual complement protease folds the numbering is as follows: C1s 438-680, C1r 464-702, MASP-1 449-696, MASP-2 445-684, C2 464-744, FB 477-757, FD 26-253, and FI 340-574. (B) PAE plots for each S12c5 model. In each plot S12c5 corresponds to residues 19-399. The residue numbering for individual complement protease models corresponds to: C1s 438-680, C1r 464-702, MASP-1 449-696, MASP-2 445-684, C2 464-744, FB 477-757, FD 26-253, and FI 340-574.

## References

1. Organization WH. Vector-borne diseases World Health Organization2020 [

2. Control of Neglected Tropical Diseases GMP, High Impact Epidemics, Vector Control and Resistance, editor Global Vector Control Response WHA 70.162017.

3. CDC. Lyme Disease Surveillance and Data 2025 [Available from: https://www.cdc.gov/lyme/data-research/facts-stats/index.html?utm_source=chatgpt.com.

4. Kugeler KJ, Schwartz AM, Delorey MJ, Mead PS, Hinckley AF. Estimating the Frequency of Lyme Disease Diagnoses, United States, 2010-2018. Emerg Infect Dis. 2021;27(2):616-9.

5. Comstedt P, Schüler W, Meinke A, Lundberg U. The novel Lyme borreliosis vaccine VLA15 shows broad protection against Borrelia species expressing six different OspA serotypes. PLoS One. 2017;12(9):e0184357.

6. Thanassi WT, Schoen RT. The Lyme disease vaccine: conception, development, and implementation. Ann Intern Med. 2000;132(8):661–8.

7. Littman MP, Gerber B, Goldstein RE, Labato MA, Lappin MR, Moore GE. ACVIM consensus update on Lyme borreliosis in dogs and cats. J Vet Intern Med. 2018;32(3):887–903.

8. Nigrovic LE, Thompson KM. The Lyme vaccine: a cautionary tale. Epidemiol Infect. 2007;135(1):1–8.

9. Trager W. Acquired Immunity to Ticks. The Journal of Parasitology. 1939;25:57–81.

10. Wikel SK. The induction of host resistance to tick infestation with a salivary gland antigen. Am J Trop Med Hyg. 1981;30(1):284–8.

11. Wikel SK, Ramachandra RN, Bergman DK, Burkot TR, Piesman J. Infestation with pathogen-free nymphs of the tick Ixodes scapularis induces host resistance to transmission of Borrelia burgdorferi by ticks. Infect Immun. 1997;65(1):335–8.

12. Nazario S, Das S, de Silva AM, Deponte K, Marcantonio N, Anderson JF, et al. Prevention of Borrelia burgdorferi transmission in guinea pigs by tick immunity. Am J Trop Med Hyg. 1998;58(6):780–5.

13. Narasimhan S, Booth CJ, Philipp MT, Fikrig E, Embers ME. Repeated Tick Infestations Impair Borrelia burgdorferi Transmission in a Non-Human Primate Model of Tick Feeding. Pathogens. 2023;12(1).

14. Narasimhan S, Deponte K, Marcantonio N, Liang X, Royce TE, Nelson KF, et al. Immunity against Ixodes scapularis salivary proteins expressed within 24 hours of attachment thwarts tick feeding and impairs Borrelia transmission. PLoS One. 2007;2(5):e451.

15. Dai J, Wang P, Adusumilli S, Booth CJ, Narasimhan S, Anguita J, Fikrig E. Antibodies against a tick protein, Salp15, protect mice from the Lyme disease agent. Cell Host Microbe. 2009;6(5):482-92.

16. Sajid A, Matias J, Arora G, Kurokawa C, DePonte K, Tang X, et al. mRNA vaccination induces tick resistance and prevents transmission of the Lyme disease agent. Sci Transl Med. 2021;13(620):eabj9827.

17. Nuttall PA, Labuda M. Tick-host interactions: saliva-activated transmission. Parasitology. 2004;129 Suppl:S177-89.

18. Severinová J, Salát J, Krocová Z, Reznícková J, Demová H, Horká H, Kopecký J. Co-inoculation of Borrelia afzelii with tick salivary gland extract influences distribution of immunocompetent cells in the skin and lymph nodes of mice. Folia Microbiol (Praha). 2005;50(5):457–63.

19. Kuthejlová M, Kopecký J, Stepánová G, Macela A. Tick salivary gland extract inhibits killing of Borrelia afzelii spirochetes by mouse macrophages. Infect Immun. 2001;69(1):575–8.

20. Zeidner NS, Schneider BS, Nuncio MS, Gern L, Piesman J. Coinoculation of Borrelia spp. with tick salivary gland lysate enhances spirochete load in mice and is tick species-specific. J Parasitol. 2002;88(6):1276–8.

21. Pechová J, Stĕpánová G, Kovár L, Kopecký J. Tick salivary gland extract-activated transmission of Borrelia afzelii spirochaetes. Folia Parasitol (Praha). 2002;49(2):153–9.

22. Kim TK, Radulovic Z, Mulenga A. Target validation of highly conserved Amblyomma americanum tick saliva serine protease inhibitor 19. Ticks Tick Borne Dis. 2016;7(3):405–14.

23. Kim TK, Tirloni L, Berger M, Diedrich JK, Yates JR, 3rd, Termignoni C, et al. Amblyomma americanum serpin 41 (AAS41) inhibits inflammation by targeting chymase and chymotrypsin. Int J Biol Macromol. 2020;156:1007-21.

24. Kim TK, Tirloni L, Bencosme-Cuevas E, Kim TH, Diedrich JK, Yates JR, 3rd, Mulenga A. Borrelia burgdorferi infection modifies protein content in saliva of Ixodes scapularis nymphs. BMC Genomics. 2021;22(1):152.

25. Tirloni L, Islam MS, Kim TK, Diedrich JK, Yates JR, 3rd, Pinto AF, et al. Saliva from nymph and adult females of Haemaphysalis longicornis: a proteomic study. Parasit Vectors. 2015;8:338.

26. Tirloni L, Reck J, Terra RM, Martins JR, Mulenga A, Sherman NE, et al. Proteomic analysis of cattle tick Rhipicephalus (Boophilus) microplus saliva: a comparison between partially and fully engorged females. PLoS One. 2014;9(4):e94831.

27. Richter D, Matuschka FR, Spielman A, Mahadevan L. How ticks get under your skin: insertion mechanics of the feeding apparatus of Ixodes ricinus ticks. Proc Biol Sci. 2013;280(1773):20131758.

28. Francischetti I, S·-Nunes A, Mans B, Santos IM, Ribeiro JMC. The role of saliva in tick feeding. Frontiers in bioscience. 2009;14:2051–88.

29. Bakshi M, Kim TK, Mulenga A. Disruption of blood meal-responsive serpins prevents Ixodes scapularis from feeding to repletion. Ticks Tick Borne Dis. 2018;9(3):506–18.

30. Bakshi M, Kim TK, Porter L, Mwangi W, Mulenga A. Amblyomma americanum ticks utilizes countervailing pro and anti-inflammatory proteins to evade host defense. PLoS Pathog. 2019;15(11):e1008128.

31. Hollmann T, Kim TK, Tirloni L, Radulović Ž M, Pinto AFM, Diedrich JK, et al. Identification and characterization of proteins in the Amblyomma americanum tick cement cone. Int J Parasitol. 2018;48(3-4):211–24.

32. Ibelli AM, Kim TK, Hill CC, Lewis LA, Bakshi M, Miller S, et al. A blood meal-induced Ixodes scapularis tick saliva serpin inhibits trypsin and thrombin, and interferes with platelet aggregation and blood clotting. Int J Parasitol. 2014;44(6):369–79.

33. Kim TK, Tirloni L, Radulovic Z, Lewis L, Bakshi M, Hill C, et al. Conserved Amblyomma americanum tick Serpin19, an inhibitor of blood clotting factors Xa and XIa, trypsin and plasmin, has anti-haemostatic functions. Int J Parasitol. 2015;45(9-10):613–27.

34. Mulenga A, Kim T, Ibelli AM. Amblyomma americanum tick saliva serine protease inhibitor 6 is a cross-class inhibitor of serine proteases and papain-like cysteine proteases that delays plasma clotting and inhibits platelet aggregation. Insect Mol Biol. 2013;22(3):306–19.

35. Yu Y, Cao J, Zhou Y, Zhang H, Zhou J. Isolation and characterization of two novel serpins from the tick Rhipicephalus haemaphysaloides. Ticks Tick Borne Dis. 2013;4(4):297–303.

36. Schuijt TJ, Hovius JW, van Burgel ND, Ramamoorthi N, Fikrig E, van Dam AP. The tick salivary protein Salp15 inhibits the killing of serum-sensitive Borrelia burgdorferi sensu lato isolates. Infect Immun. 2008;76(7):2888–94.

37. Denisov SS, Heinzmann ACA, Vajen T, Vries MHM, Megens RTA, Suylen D, et al. Tick Saliva Protein Evasin-3 Allows for Visualization of Inflammation in Arteries through Interactions with CXC-Type Chemokines Deposited on Activated Endothelium. Bioconjug Chem. 2020;31(3):948–55.

38. Mihaljica D, Marković D, Radulović Ž, Mulenga A, Ćakić S, Sukara R, et al. Ixodes ricinus immunogenic saliva protein, homologue to Amblyomma americanum AV422: Determining its potential for use in tick bite confirmation. Ticks Tick Borne Dis. 2017;8(3):391–5.

39. Zeng YT, Liu YN, Chen ZH, Chen Q, Sun KP. The impact of antithrombin III supplementation on prognosis during extracorporeal membrane oxygenation: a systematic review and meta-analysis. Ann Med. 2025;57(1):2542439.

40. Olson ST, Gettins PG. Regulation of proteases by protein inhibitors of the serpin superfamily. Prog Mol Biol Transl Sci. 2011;99:185–240.

41. Mulenga A, Sugino M, Nakajim M, Sugimoto C, Onuma M. Tick-Encoded serine proteinase inhibitors (serpins); potential target antigens for tick vaccine development. J Vet Med Sci. 2001;63(10):1063–9.

42. Porter L, Radulović Ž, Kim T, Braz GR, Da Silva Vaz I, Jr., Mulenga A. Bioinformatic analyses of male and female Amblyomma americanum tick expressed serine protease inhibitors (serpins). Ticks Tick Borne Dis. 2015;6(1):16–30.

43. Mudenda L, Pierlé SA, Turse JE, Scoles GA, Purvine SO, Nicora CD, et al. Proteomics informed by transcriptomics identifies novel secreted proteins in Dermacentor andersoni saliva. Int J Parasitol. 2014;44(13):1029–37.

44. Kim TK, Tirloni L, Pinto AF, Moresco J, Yates JR, 3rd, da Silva Vaz I, Jr., Mulenga A. Ixodes scapularis Tick Saliva Proteins Sequentially Secreted Every 24 h during Blood Feeding. PLoS Negl Trop Dis. 2016;10(1):e0004323.

45. Tirloni L, Kim TK, Pinto AFM, Yates JR, 3rd, da Silva Vaz I, Jr., Mulenga A. Tick-Host Range Adaptation: Changes in Protein Profiles in Unfed Adult Ixodes scapularis and Amblyomma americanum Saliva Stimulated to Feed on Different Hosts. Front Cell Infect Microbiol. 2017;7:517.

46. Radulović Z, Porter LM, Kim TK, Mulenga A. Comparative bioinformatics, temporal and spatial expression analyses of Ixodes scapularis organic anion transporting polypeptides. Ticks Tick Borne Dis. 2014;5(3):287–98.

47. Nguyen TT, Kim TH, Bencosme-Cuevas E, Berry J, Gaithuma ASK, Ansari MA, et al. A tick saliva serpin, IxsS17 inhibits host innate immune system proteases and enhances host colonization by Lyme disease agent. PLoS Pathog. 2024;20(2):e1012032.

48. Tirloni L, Kim TK, Berger M, Termignoni C, da Silva Vaz I, Jr., Mulenga A. Amblyomma americanum serpin 27 (AAS27) is a tick salivary anti-inflammatory protein secreted into the host during feeding. PLoS Negl Trop Dis. 2019;13(8):e0007660.

49. Bencosme-Cuevas E, Kim TK, Nguyen TT, Berry JR, Li J, Adams LG, et al. Ixodes scapularis nymph saliva protein blocks host inflammation and complement-mediated killing of Lyme disease agent, Borrelia burgdorferi. Frontiers in Cellular and Infection Microbiology. 2023;13.

50. Nuss AB, Lomas JS, Reyes JB, Garcia-Cruz O, Lei W, Sharma A, et al. The highly improved genome of Ixodes scapularis with X and Y pseudochromosomes. Life Sci Alliance. 2023;6(12).

51. Mulenga A, Khumthong R, Chalaire KC. Ixodes scapularis tick serine proteinase inhibitor (serpin) gene family; annotation and transcriptional analysis. BMC Genomics. 2009;10:217.

52. Alex Samuel Kiarie Gaithuma T-TN, Albert Mulenga. Genes 2026; In Press. Chromosome-scale atlas of Ixodes scapularis serine protease inhibitors. Genes. 2026.

53. Sekine H, Takahashi M, Iwaki D, Fujita T. The role of MASP-1/3 in complement activation. Adv Exp Med Biol. 2013;735:41–53.

54. Dobó J, Kocsis A, Farkas B, Demeter F, Cervenak L, Gál P. The Lectin Pathway of the Complement System-Activation, Regulation, Disease Connections and Interplay with Other (Proteolytic) Systems. Int J Mol Sci. 2024;25(3).

55. Horvath AJ, Lu BG, Pike RN, Bottomley SP. Methods to measure the kinetics of protease inhibition by serpins. Methods Enzymol. 2011;501:223–35.

56. Saleem A, Blifeld C, Saleh SA, Yawn DH, Mace ML, Schwartz M, Crawford ES. Viscoelastic measurement of clot formation: a new test of platelet function. Ann Clin Lab Sci. 1983;13(2):115–24.

57. Evrard J, Siriez R, Morimont L, Thémans P, Laloy J, Bouvy C, et al. Optimal wavelength for the clot waveform analysis: Determination of the best resolution with minimal interference of the reagents. Int J Lab Hematol. 2019;41(3):316–24.

58. Pieters M, Guthold M, Nunes CM, de Lange Z. Interpretation and Validation of Maximum Absorbance Data Obtained from Turbidimetry Analysis of Plasma Clots. Thromb Haemost. 2020;120(1):44–54.

59. Lacroix MB, Aude CA, Arlaud GJ, Colomb MG. Isolation and functional characterization of the proenzyme form of the catalytic domains of human C1r. Biochem J. 1989;257(3):885–91.

60. Gaboriaud C, Rossi V, Bally I, Arlaud GJ, Fontecilla-Camps JC. Crystal structure of the catalytic domain of human complement c1s: a serine protease with a handle. Embo j. 2000;19(8):1755–65.

61. Heise CT, Nicholls JR, Leamy CE, Wallis R. Impaired secretion of rat mannose-binding protein resulting from mutations in the collagen-like domain. J Immunol. 2000;165(3):1403–9.

62. Matsushita M, Thiel S, Jensenius JC, Terai I, Fujita T. Proteolytic activities of two types of mannose-binding lectin-associated serine protease. J Immunol. 2000;165(5):2637–42.

63. Milacic M, Beavers D, Conley P, Gong C, Gillespie M, Griss J, et al. The Reactome Pathway Knowledgebase 2024. Nucleic Acids Res. 2024;52(D1):D672–d8.

64. Karnaukhova E. C1-Inhibitor: Structure, Functional Diversity and Therapeutic Development. Curr Med Chem. 2022;29(3):467–88.

65. Lord MS, Melrose J, Day AJ, Whitelock JM. The Inter-α-Trypsin Inhibitor Family: Versatile Molecules in Biology and Pathology. J Histochem Cytochem. 2020;68(12):907–27.

66. Rezaie AR, Giri H. Anticoagulant and signaling functions of antithrombin. J Thromb Haemost. 2020;18(12):3142–53.

67. Kakoki M, Smithies O. The kallikrein-kinin system in health and in diseases of the kidney. Kidney Int. 2009;75(10):1019–30.

68. Garrigues RJ, Garrison MP, Garcia BL. The Crystal Structure of the Michaelis-Menten Complex of C1 Esterase Inhibitor and C1s Reveals Novel Insights into Complement Regulation. J Immunol. 2024;213(5):718–29.

69. Booth CE, Jr., Powell-Pierce AD, Skare JT, Garcia BL. Borrelia miyamotoi FbpA and FbpB Are Immunomodulatory Outer Surface Lipoproteins With Distinct Structures and Functions. Front Immunol. 2022;13:886733.

70. Labandeira-Rey M, Seshu J, Skare JT. The absence of linear plasmid 25 or 28-1 of Borrelia burgdorferi dramatically alters the kinetics of experimental infection via distinct mechanisms. Infect Immun. 2003;71(8):4608–13.

71. Labandeira-Rey M, Skare JT. Decreased infectivity in Borrelia burgdorferi strain B31 is associated with loss of linear plasmid 25 or 28-1. Infect Immun. 2001;69(1):446–55.

72. van Dam AP, Kuiper H, Vos K, Widjojokusumo A, de Jongh BM, Spanjaard L, et al. Different genospecies of Borrelia burgdorferi are associated with distinct clinical manifestations of Lyme borreliosis. Clin Infect Dis. 1993;17(4):708–17.

73. Gulia-Nuss M, Nuss AB, Meyer JM, Sonenshine DE, Roe RM, Waterhouse RM, et al. Genomic insights into the Ixodes scapularis tick vector of Lyme disease. Nat Commun. 2016;7:10507.

74. Li W, Johnson DJ, Esmon CT, Huntington JA. Structure of the antithrombin-thrombin-heparin ternary complex reveals the antithrombotic mechanism of heparin. Nat Struct Mol Biol. 2004;11(9):857–62.

75. Tollefsen DM. Vascular dermatan sulfate and heparin cofactor II. Prog Mol Biol Transl Sci. 2010;93:351–72.

76. Chantelle M. Rein URD, Frank C. Church,. Serpin–Glycosaminoglycan Interactions,. 2011. In: Methods in Enzymology [Internet]. Academic Press; [105-37].

77. Sobczak AIS, Pitt SJ, Stewart AJ. Glycosaminoglycan Neutralization in Coagulation Control. Arterioscler Thromb Vasc Biol. 2018;38(6):1258–70.

78. Cottrell GS, Amadesi S, Grady EF, Bunnett NW. Trypsin IV, a novel agonist of protease-activated receptors 2 and 4. J Biol Chem. 2004;279(14):13532–9.

79. Knecht W, Cottrell GS, Amadesi S, Mohlin J, Skåregärde A, Gedda K, et al. Trypsin IV or mesotrypsin and p23 cleave protease-activated receptors 1 and 2 to induce inflammation and hyperalgesia. J Biol Chem. 2007;282(36):26089–100.

80. Booth NA. Fibrinolysis and thrombosis. Baillieres Best Pract Res Clin Haematol. 1999;12(3):423–33.

81. Syrovets T, Lunov O, Simmet T. Plasmin as a proinflammatory cell activator. J Leukoc Biol. 2012;92(3):509–19.

82. Francischetti IM, Sa-Nunes A, Mans BJ, Santos IM, Ribeiro JM. The role of saliva in tick feeding. Front Biosci (Landmark Ed). 2009;14:2051–88.

83. Leavell KJ, Peterson MW, Gross TJ. The role of fibrin degradation products in neutrophil recruitment to the lung. Am J Respir Cell Mol Biol. 1996;14(1):53–60.

84. Gross TJ, Leavell KJ, Peterson MW. CD11b/CD18 mediates the neutrophil chemotactic activity of fibrin degradation product D domain. Thromb Haemost. 1997;77(5):894–900.

85. Menten-Dedoyart C, Faccinetto C, Golovchenko M, Dupiereux I, Van Lerberghe PB, Dubois S, et al. Neutrophil extracellular traps entrap and kill Borrelia burgdorferi sensu stricto spirochetes and are not affected by Ixodes ricinus tick saliva. J Immunol. 2012;189(11):5393–401.

86. Koenigs A, Hammerschmidt C, Jutras BL, Pogoryelov D, Barthel D, Skerka C, et al. BBA70 of Borrelia burgdorferi is a novel plasminogen-binding protein. J Biol Chem. 2013;288(35):25229–43.

87. Fuchs H, Wallich R, Simon MM, Kramer MD. The outer surface protein A of the spirochete Borrelia burgdorferi is a plasmin(ogen) receptor. Proc Natl Acad Sci U S A. 1994;91(26):12594–8.

88. Önder Ö, Humphrey PT, McOmber B, Korobova F, Francella N, Greenbaum DC, Brisson D. OspC is potent plasminogen receptor on surface of Borrelia burgdorferi. J Biol Chem. 2012;287(20):16860–8.

89. Brouwer MAE, van de Schoor FR, Vrijmoeth HD, Netea MG, Joosten LAB. A joint effort: The interplay between the innate and the adaptive immune system in Lyme arthritis. Immunol Rev. 2020;294(1):63–79.

90. Coleman JL, Roemer EJ, Benach JL. Plasmin-coated borrelia Burgdorferi degrades soluble and insoluble components of the mammalian extracellular matrix. Infect Immun. 1999;67(8):3929–36.

91. Itoh Y. Membrane-type matrix metalloproteinases: Their functions and regulations. Matrix Biol. 2015;44–46:207-23.

92. Šimo L, Kazimirova M, Richardson J, Bonnet SI. The Essential Role of Tick Salivary Glands and Saliva in Tick Feeding and Pathogen Transmission. Front Cell Infect Microbiol. 2017;7:281.

93. Valenzuela JG, Charlab R, Mather TN, Ribeiro JM. Purification, cloning, and expression of a novel salivary anticomplement protein from the tick, Ixodes scapularis. J Biol Chem. 2000;275(25):18717–23.

94. Denisov SS, Ippel JH, Castoldi E, Mans BJ, Hackeng TM, Dijkgraaf I. Molecular basis of anticoagulant and anticomplement activity of the tick salivary protein Salp14 and its homologs. J Biol Chem. 2021;297(1):100865.

95. Tyson K, Elkins C, Patterson H, Fikrig E, de Silva A. Biochemical and functional characterization of Salp20, an Ixodes scapularis tick salivary protein that inhibits the complement pathway. Insect Mol Biol. 2007;16(4):469–79.

96. Bencosme-Cuevas E. KT, Nguyen TT, Berry JR., Li J., Adams LG., Smith LA., Swale DR., Kaufmann SHE, Jones-Hall H., Mulenga A. Ixodes scapularis nymph saliva protein blocks host inflammation and complement-mediated killing of Lyme disease agent, Borrelia burgdorferi. Frontiers in Cellular and Infection Microbiology. 2023;13.

97. Qian WJ, Kaleta DT, Petritis BO, Jiang H, Liu T, Zhang X, et al. Enhanced detection of low abundance human plasma proteins using a tandem IgY12-SuperMix immunoaffinity separation strategy. Mol Cell Proteomics. 2008;7(10):1963–73.

98. Zubarev RA. The challenge of the proteome dynamic range and its implications for in-depth proteomics. Proteomics. 2013;13(5):723–6.

99. Thiel S, Jensen L, Degn SE, Nielsen HJ, Gál P, Dobó J, Jensenius JC. Mannan-binding lectin (MBL)-associated serine protease-1 (MASP-1), a serine protease associated with humoral pattern-recognition molecules: normal and acute-phase levels in serum and stoichiometry of lectin pathway components. Clin Exp Immunol. 2012;169(1):38–48.

100. Frauenknecht V, Thiel S, Storm L, Meier N, Arnold M, Schmid JP, et al. Plasma levels of mannan-binding lectin (MBL)-associated serine proteases (MASPs) and MBL-associated protein in cardio- and cerebrovascular diseases. Clin Exp Immunol. 2013;173(1):112–20.

101. Venkatraman Girija U, Gingras AR, Marshall JE, Panchal R, Sheikh MA, Harper JA, et al. Structural basis of the C1q/C1s interaction and its central role in assembly of the C1 complex of complement activation. Proc Natl Acad Sci U S A. 2013;110(34):13916–20.

102. Sullivan KE. The yin and the yang of early classical pathway complement disorders. Clin Exp Immunol. 2022;209(2):151–60.

103. Hourcade DE. The role of properdin in the assembly of the alternative pathway C3 convertases of complement. J Biol Chem. 2006;281(4):2128–32.

104. Blatt AZ, Pathan S, Ferreira VP. Properdin: a tightly regulated critical inflammatory modulator. Immunol Rev. 2016;274(1):172–90.

105. Pangburn MK, Schreiber RD, Müller-Eberhard HJ. Human complement C3b inactivator: isolation, characterization, and demonstration of an absolute requirement for the serum protein beta1H for cleavage of C3b and C4b in solution. J Exp Med. 1977;146(1):257–70.

106. Liao M, Zhou J, Gong H, Boldbaatar D, Shirafuji R, Battur B, et al. Hemalin, a thrombin inhibitor isolated from a midgut cDNA library from the hard tick Haemaphysalis longicornis. J Insect Physiol. 2009;55(2):164–73.

107. Gao X, Shi L, Zhou Y, Cao J, Zhang H, Zhou J. Characterization of the anticoagulant protein Rhipilin-1 from the Rhipicephalus haemaphysaloides tick. J Insect Physiol. 2011;57(2):339–43.

108. Esmon CT. Regulation of blood coagulation. Biochim Biophys Acta. 2000;1477(1-2):349–60.

109. Sajanti EM, Gröndahl-Yli-Hannuksela K, Kauko T, He Q, Hytönen J. Lyme Borreliosis and Deficient Mannose-Binding Lectin Pathway of Complement. J Immunol. 2015;194(1):358–63.

110. Coumou J, Wagemakers A, Narasimhan S, Schuijt TJ, Ersoz JI, Oei A, et al. The role of Mannose Binding Lectin in the immune response against Borrelia burgdorferi sensu lato. Sci Rep. 2019;9(1):1431.

111. Mekaj YH, Mekaj AY, Duci SB, Miftari EI. New oral anticoagulants: their advantages and disadvantages compared with vitamin K antagonists in the prevention and treatment of patients with thromboembolic events. Ther Clin Risk Manag. 2015;11:967–77.

112. Esmon CT. Targeting factor Xa and thrombin: impact on coagulation and beyond. Thromb Haemost. 2014;111(4):625–33.

113. Radulović Ž M, Mulenga A. Heparan sulfate/heparin glycosaminoglycan binding alters inhibitory profile and enhances anticoagulant function of conserved Amblyomma americanum tick saliva serpin 19. Insect Biochem Mol Biol. 2017;80:1–10.

114. Ansari MA, Nguyen TT, Kocurek KI, Kim WTH, Kim TK, Mulenga A. Recombinant Ixodes scapularis Calreticulin Binds Complement Proteins but Does Not Protect Borrelia burgdorferi from Complement Killing. Pathogens. 2024;13(7).

115. Abramson J, Adler J, Dunger J, Evans R, Green T, Pritzel A, et al. Accurate structure prediction of biomolecular interactions with AlphaFold 3. Nature. 2024;630(8016):493–500.

116. Kurtenbach K, Sewell HS, Ogden NH, Randolph SE, Nuttall PA. Serum complement sensitivity as a key factor in Lyme disease ecology. Infect Immun. 1998;66(3):1248–51.

117. Skare JT, Garcia BL. Complement Evasion by Lyme Disease Spirochetes. Trends Microbiol. 2020.

118. Xie J, Zhi H, Garrigues RJ, Keightley A, Garcia BL, Skare JT. Structural determination of the complement inhibitory domain of Borrelia burgdorferi BBK32 provides insight into classical pathway complement evasion by Lyme disease spirochetes. PLoS Pathog. 2019;15(3):e1007659.

119. Garcia BL, Zhi H, Wager B, Höök M, Skare JT. Borrelia burgdorferi BBK32 Inhibits the Classical Pathway by Blocking Activation of the C1 Complement Complex. PLoS Pathog. 2016;12(1):e1005404.

120. Garrigues RJ, Powell-Pierce AD, Hammel M, Skare JT, Garcia BL. A Structural Basis for Inhibition of the Complement Initiator Protease C1r by Lyme Disease Spirochetes. J Immunol. 2021;207(11):2856–67.

121. Van Laar TA, Hole C, Rajasekhar Karna SL, Miller CL, Reddick R, Wormley FL, Seshu J. Statins reduce spirochetal burden and modulate immune responses in the C3H/HeN mouse model of Lyme disease. Microbes Infect. 2016;18(6):430–5.

